# Evaluation of *P. aeruginosa* attachment on mineralized collagen scaffolds and addition of manuka honey to increase mesenchymal stem cell osteogenesis

**DOI:** 10.1101/2022.01.28.478244

**Authors:** Marley J. Dewey, Alan J. Collins, Aleczandria Tiffany, Victoria R. Barnhouse, Crislyn Lu, Vasiliki Kolliopoulos, Noreen J. Hickok, Brendan A.C. Harley

## Abstract

The design of biomaterials to regenerate bone is likely to increasingly require modifications that reduce bacterial attachment and biofilm formation as infection during wound regeneration can significantly impede tissue repair and typically requires surgical intervention to restart the healing process. Here, we investigate the ability of a mineralized collagen biomaterial to natively resist infection as well as how the addition of manuka honey affects bacterial colonization and mesenchymal stem cell osteogenesis. We incorporate manuka honey into these scaffolds via either direct fabrication into the scaffold microarchitecture or via soaking the scaffold in a solution of Manuka honey after fabrication. Direct incorporation results in a change in the surface characteristics and porosity of mineralized collagen scaffolds. Soaking scaffolds in honey concentrations greater than 10% had significant negative effects on mesenchymal stem cell metabolic activity but soaking or incorporating 5% honey had no impact on endothelial cell tube formation. Soaking and incorporating 5% honey into scaffolds reduced metabolic activity of mesenchymal stem cells, however, soaking 5% honey into scaffolds increased calcium and phosphorous mineral formation, osteoprotegerin release, and alkaline phosphatase activity. The addition of manuka honey did not prevent *P. aeruginosa* attachment but may be able to limit attachment of other common wound-colonizing bacteria. Overall, our results demonstrate the potential for soaking mineralized collagen scaffolds in 5% manuka honey to increase osteogenesis of mesenchymal stem cells.

## 1. Introduction

Craniomaxillofacial (CMF) defects occur at all ages; and encompass cleft palate birth defects traumatic injuries, cancer resection and bone loss from dentures [1–4]. Characteristically, these defects encompass large portions of bone from the skull or jaw and require a bone replacement to regenerate this missing tissue. Autografts have the highest success rates for complete bone regeneration but involve secondary surgeries with pain and increased risk of bone morbidity at the site(s) of harvest [4, 5]. To circumvent the problems associated with a limited supply of autografts as well as the complications associated with its harvesting, interest has turned to the development of allogenic implantable biomaterials. However, due to the size of the defects among other obstacles, no biomaterial strategy has been able to match the success of autografts. A dire complication regardless of treatment strategy is infection. Treatment requires aggressive use of antimicrobials and at least one additional operation to remove the implant. If the infection can be cleared, then the implantation process can be re-started, albeit with an increased risk for infection; unsuccessful treatments result in disfigurement and even death.

Implanted materials account for 45% of all hospital-contracted infections, and there is a drastic and rapidly rising cost to treating these, with $150-200 million spent in the US in 1993, and $11 billion spent in the US annually in 2001 [6–8]. For craniomaxillofacial defects, infections occur in as many as 40% of patients with biomaterial implants [9–12]. The highest rates are associated with trauma and battlefield injuries, where wound contamination is common [13]. Current biomaterials seeking to regenerate CMF defects have been hindered by localized infections in the implanted biomaterial. As early as 7 days post-implantation, infection was detected after CMF reconstruction using PEEK (polymer) implants, where infection rates approached 28% and resulted in complications including abscess formation, pain and swelling, osteomyelitis, and fistula formation [14]. An additional complication arises as antibiotics are the mainstay of anti-infective/prophylactic treatments [15]. However, biomaterial-adherent and biofilm bacteria are notably tolerant to antibiotics, requiring levels 100-1000X higher than the strains minimal inhibitory concentration (MIC) [16].

*Pseudomonas aeruginosa* is a gram-negative bacterium that commonly infects wound sites. *P. aeruginosa* infections can be recalcitrant to treatment due to the increasing prevalence of antibiotic resistant strains and the formation of biofilms that increase tolerance of antibiotic and chemical treatments [17]. Although *P. aeruginosa* infections in bone joint repair are uncommon (3% of 90 cases), treatment of these infections is especially difficult, and can lead to chronic osteomyelitis, or bone inflammation [17, 18]. Chronic osteomyelitis can lead to pain at the injury site, bone deformations, disrupted vascular networks, and limited mobility [18]. Of concern, use of low doses of antibiotics for long-term infection prevention can promote biofilm formation and tolerance to antibiotics [19]. Additionally, use of treatments such as tumor necrosis factor alpha (TNF-α) for autoimmune disorders, have the possibility to increase the risk of *P. aeruginosa* infection. In one case, a dormant infection at the site of an old shrapnel wound was re-activated following TNF-α treatment, leading to osteomyelitis and soft-tissue abscess [20]. Furthermore, CMF defects often have persistent inflammation due to the size of missing tissue, and inflamed tissue can block the ability of antibiotics to reach the infected site [15].

The growing problem of antimicrobial resistance and the inability of antibiotics to penetrate biofilms in wounds suggests opportunities to develop biomaterials that may alter bacterial attachment. One intriguing option is the use of naturally occurring substances which might extend resistance to biofilms. Manuka honey has been investigated recently as an alternative to antibiotics due to known antibacterial properties. Notably, leptospermum (manuka) honeys have demonstrated efficacy against a wide range of both gram-negative and -positive bacteria [21], including bactericidal properties against both *S. aureus* and *Pseudomonas aeruginosa* [22]. The antibacterial and wound-healing properties of honeys has been attributed to its high sugar content, low pH, hydrogen peroxide and enzyme content, as well as high levels of methylglyoxal (MGO) [21, 23]. Critically, there has been no data to indicate bacteria developed resistance to manuka honeys [21, 23, 24], and manuka honey has been incorporated into electrospun scaffolds, hydrogels, and cryogels [23, 25, 26]. While manuka honey has demonstrated prevention of bacterial attachment, few studies have examined the details of how honey may affect mesenchymal stem cell activity and regenerative potential.

Mineralized collagen biomaterials have been researched extensively for CMF defect repair and have demonstrated excellent *in vitro* biocompatibility and mesenchymal stem cell osteogenesis, as well as bone regeneration *in vivo* [27–43]. While growth potential of bacteria in these scaffolds is not well known, it is likely that bacteria that commonly infect bone wounds, such as *Staphylococcus aureus* and *Pseudomonas aeruginosa*, will be able to colonize collagen-based biomaterials [44, 45]. Furthermore, type I collagen has been shown to enhance attachment of bacteria such as *Streptococcus mutans* to dentin [46]. Hence, the goal of this project was to incorporate manuka honey into mineralized collagen scaffolds and define the influence of honey incorporation on bacteria attachment. We report the effect of manuka honey in a class of mineralized collagen scaffolds towards preventing *P. aeruginosa* attachment, a bacterium that readily forms antibiotic-tolerant biofilms [47], and the effect of manuka honey on osteogenesis of mesenchymal stem cells on these scaffolds. Additionally, as vascularization plays an important role in regenerative healing, we examine the influence of honey concentrations on endothelial tube formation. We report two methods of incorporating manuka honey into mineralized collagen scaffolds, direct incorporation of honey into the collagen scaffold during fabrication (*honey incorporated*) versus soaking mineralized scaffolds in a solution of manuka honey (*honey soaked*). We subsequently report the metabolic activity, mineral formation, MSC-mediated secretion of osteoprotegerin (OPG, a potent osteoclast inhibitory glycoprotein) release, and alkaline phosphatase activity of human mesenchymal stem cells (hMSCs). We then assess *P. aeruginosa* attachment and proliferation within the scaffolds and examine shifts in endothelial cell tube formation capacity in response to the conditioned media generated by hMSCs cultured within mineralized collagen scaffolds containing manuka honey. These studies provide key information regarding the role of Manuka honey functionalized scaffolds on biomarkers of MSC osteogenesis and *P. aeruginosa* colonization.

## 2. Materials and Methods

### 2.1 Fabrication of mineralized collagen scaffolds with manuka honey

Two versions of manuka honey-containing mineralized collagen scaffolds were fabricated, via soaking or during homogenization (**Fig. 1**). Mineralized collagen scaffolds were fabricated following previous studies [30, 48, 49]. Briefly, 1.9 w/v% type I bovine collagen (Sigma-Aldrich, Missouri, USA) was homogenized together with a 40 w/v% mineral solution comprising phosphoric acid (Fisher Scientific, New Hampshire, USA) and calcium hydroxide (Sigma Aldrich). After blending these together, 0.84 v/v% chondroitin-6-sulfate (Sigma-Aldrich) and calcium nitrate tetrahydrate (Sigma Aldrich) were added [29, 30, 48, 50]. The suspension was refrigerated overnight and then 24 mL of this suspension was added to a 75 × 75 mm aluminum pan for *in vitro* testing and 1.5 mL to 10 mm dia. × 10 mm height polysulfone molds for mechanical compression testing. These molds were then freeze-dried using a Genesis freeze-dryer (VirTis, New York, USA) by a constant decrease in temperature at a rate of 1°C/min until reaching and holding at −10°C for 2 hours. After completion, these were brought to room temperature and pressure and stored in a desiccator or at 4°C until use. These scaffolds are referred to as 0 v/v% manuka honey containing scaffolds. To create scaffolds for porosity, cell and bacterial cultures, biopsy punches (12 mm dia. and 6 mm dia.) were used on the resulting mineralized collagen sheet formed from the aluminum pan.

**Fig. 1.**
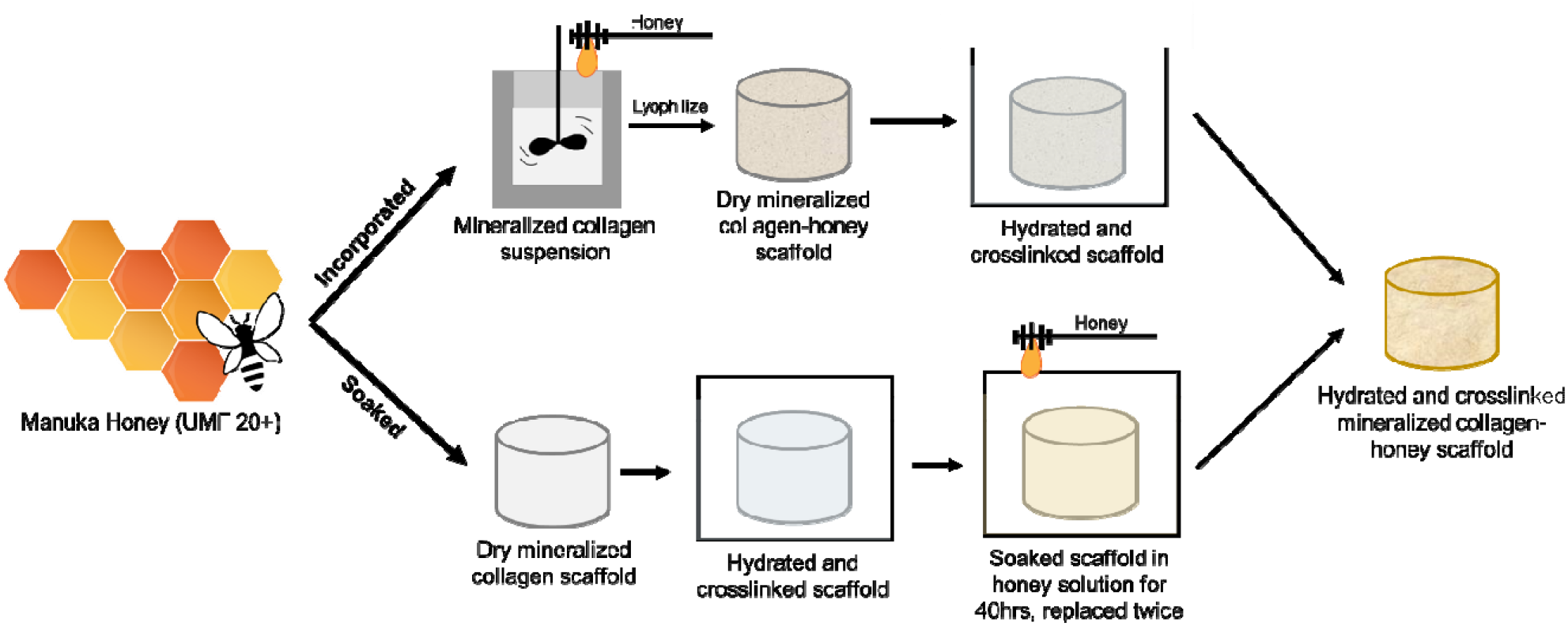
Fabrication of manuka honey-mineralized collagen scaffolds by method of incorporation or soaking. To fabricate incorporated manuka honey scaffolds, the desired v/v% manuka honey was added to a complete mineralized collagen suspension and blended together with a homogenizer. Suspension was then lyophilized to create dry honey incorporated mineralized collagen scaffolds. Scaffolds were then sterilized and hydrated and crosslinked before any *in vitro* testing. To fabricate soaked manuka honey scaffolds, normal dry mineralized collagen scaffolds were sterilized, hydrated, and crosslinked, and then soaked in solution containing the desired v/v% manuka honey for 40 hours with a total of two replacements of honey-containing solution.

#### Honey incorporated scaffolds

To create 2 and 5 v/v% manuka honey incorporated mineralized collagen scaffolds (hereafter referred to as 2% and 5%), manuka honey was added to mineralized collagen suspension during the blending process. The appropriate volume of honey (depending on final concentration in suspension) was added to 4 mL of DI water and mixed thoroughly before slowly adding to a completed mineralized collagen suspension while blending. The mineralized collagen-honey incorporated suspension contained all listed components in the above section plus 2 or 5 v/v% manuka honey. All honey used with mineralized collagen scaffolds was of the brand Australia’s Manuka P/L (Good Natured Inc, Vancouver, BC Canada) measuring MGO 820+ and NPA 20+. Scaffolds were then fabricated in a similar manner as above, by adding 24 mL of the honey-mineralized collagen suspension to aluminum pans or polysulfone molds and freeze drying at the same conditions.

#### Honey soaked scaffolds

To create various (2-50) v/v% manuka honey soaked mineralized collagen scaffolds, honey was added to these after the lyophilization process. Mineralized collagen scaffolds were first sterilized, hydrated in PBS, then crosslinked (detailed in later sections), before soaking in either cell culture medium or PBS containing a specific v/v% manuka honey for approximately 40 hrs with two changes of fresh medium containing manuka honey. Manuka honey solutions were always made fresh day-of use.

### 2.2 Porosity

The porosity of mineralized collagen honey incorporated scaffolds was measured via soaking in isopropanol and measuring uptake of this solution into the scaffolds [51]. Dry 12 mm biopsy punches of 2% and 5% honey incorporated scaffolds were compared to mineralized collagen scaffolds without honey (n=8). The initial volume and weight were measured before soaking for 24 hrs in isopropanol at room temperature on a shaker. After this, scaffolds were briefly dried for 1 minute on a filter paper before measuring the new weight. The following equation was then used to calculate the porosity and swelling of the scaffolds [52].

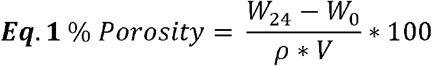

Where W_24_ represents the weight of the scaffolds after 24 hrs, W_0_ represents the initial scaffold weight, ρ represents the density of isopropanol, and V represents the initial volume of the scaffold.

### 2.3 Sterilization, hydration, and crosslinking of scaffolds

Before *in vitro* tests, scaffolds were sterilized via ethylene oxide for 12 hours with an AN74i Anprolene gas sterilizer (Andersen Sterilizers Inc., Haw River, NC). After sterilization, scaffolds were handled following sterile procedures in a biosafety cabinet. Scaffolds were hydrated following previously described procedures [49, 50, 53–57]. Briefly, scaffolds were hydrated in 100% ethanol, then in multiple washes with PBS, and then crosslinked with carbodiimide chemistry in PBS solution:1-ethyl-3-(3-dimethylaminopropyl) carbodiimide hydrochloride (EDC, Sigma-Aldrich) and N-hydroxysuccinimide (NHS, Sigma-Aldrich) at a molar ration of 5:2:1 EDAC:NHS:COOH, with carboxylic acid groups present on the collagen backbone. After crosslinking, scaffolds were washed again in PBS and then soaked in the appropriate solution. Mineralized collagen and mineralized collagen honey incorporated scaffolds were soaked in basal cell culture media (for bacterial and cell studies) or PBS (for release, pH, and mechanical testing). Mineralized collagen honey soaked scaffolds were added to either solutions of cell culture media or PBS containing the desired v/v% manuka honey.

### 2.4 Glucose and methylglyoxal release from scaffolds

To measure the amount of honey released from scaffolds, glucose and MGO released from the scaffolds was quantified. Hydrated and crosslinked mineralized collagen scaffolds without honey and 2% and 5% honey incorporated and honey soaked scaffolds measuring 6 mm in diameter were added to a 24-well plate in 1 mL of PBS per well on a shaker at 37°C. PBS was collected and replaced at days 1, 4, 7, and 14. A glucose release curve was created by evaluating the glucose released into PBS at days 1, 4, 7, and 14 with a fluorometric glucose assay kit (Cell Biolabs Inc, California, USA). An MGO assay (methylglyoxal assay kit, BioVision, California, USA) was used on only the 0% honey, 5% honey incorporated, and 5% honey soaked scaffolds after 1 day of soaking in PBS. No dilution of samples was necessary and an n=6 sample group size was used.

### 2.5 pH testing

Hydrated and crosslinked mineralized collagen scaffolds without honey and 2% and 5% honey incorporated and honey soaked scaffolds measuring 6 mm in diameter were added to a 24-well plate containing 1 mL of PBS. These were added to a shaker at 37°C and the pH was measured at days 1-7, 10, and 14, replacing every day after measurement. The pH of the PBS stock solution was measured as a control (n=6).

### 2.6 Mechanical compression testing

Hydrated and crosslinked mineralized collagen scaffolds (10 mm dia. × 10 mm height) without honey and 2% and 5% honey incorporated scaffolds were mechanically compressed while wet. Scaffolds were compressed at a rate of 1 mm/min using a 5N load cell and Instron 5943 (Instron, Norwood, MA). The slope of the linear elastic portion of the stress-strain curves was analyzed to yield young’s modulus following similar protocols for open-porous scaffolds [48, 58, 59] (n=8).

### 2.7 Human mesenchymal stem cell culture

Two donors of human mesenchymal stem cells (hMSCs) at passage 5-6 were seeded on mineralized collagen scaffolds for osteogenesis experiments (seeded separately, BM-17, Lonza, Maryland, USA or RoosterBio, Maryland, USA). Before culture on scaffolds, media surrounding cells was assessed for mycoplasma contamination using a MycoAlert™ Mycoplasma Detection Kit (Lonza). All cells used on scaffolds tested negative for mycoplasma. Hydrated and crosslinked scaffolds were seeded with 100,000 total hMSCs in ultra-low attachment 24-well plate (Corning, New York, USA). First, 10 μL containing 50,000 cells was pipetted directly on one side of the scaffold for 30 min, then 10 μL of 50,000 cells was pipetted on the other side for 1.5 hours before adding basal (either phenol-containing or phenol-red free) cell culture media (low glucose DMEM and L-glutamine, 10% mesenchymal stem cell fetal bovine serum, and 1% antibiotic-antimycotic). Scaffolds with cells were then added to incubators maintaining 5% CO_2_ and 37°C with fresh media changes every 3 days.

### 2.8 Metabolic activity of hMSCs on scaffolds

To measure the cell health of hMSCs on scaffolds with honey, an AlamarBlue® assay was used (n=6). Prior to seeding hMSCs on scaffolds, a standard curve of known cell number was generated from the amount of resazurin converted to fluorescent resorufin by cells. This was measured using a fluorescent spectrophotometer (Tecan, Switzerland). To quantify metabolic activity of cells in scaffolds, scaffolds were added to a solution of AlmarBlue® (ThermoFisher Scientific, Massachusetts, USA) and media and soaked for 1.5 hours on a shaker in an incubator before reading the fluorescence of the solution. Scaffolds were then added back to original wells in the incubator. The standard curve was used to convert fluorescent readings to initial cell activity, with a value of 1 representing the cell health of the 100,000 cells seeded onto the scaffolds.

### 2.9 Mineral analysis of scaffolds after hMSC culture

The amount of calcium and phosphorous produced by hMSCs seeded onto scaffolds after 14 days of culture was assessed by inductively coupled plasma mass spectroscopy (ICP, n=6). After 14 days of hMSC culture on scaffolds, scaffolds were added to Formal-Fixx (10% neutral buffered formalin, ThermoFisher Scientific) for 24 hours at 4°C. After fixing, scaffolds were washed three times for 5 min in PBS before drying briefly on a KimWipe® and storing at −80°C. Scaffolds were dried by lyophilizing via the procedure outlined in **Section 2.1**. Inductively Coupled Plasma (ICP) Optical Emission spectrometry was then performed on fixed and dry collagen scaffolds to assess calcium and phosphorous percent mineral formation. The mass of samples was recorded, and then dissolved in concentrated nitric acid (Trace Metal Grade concentrated HNO_3_, Thermo Fischer Scientific 67-70%). After dissolving, samples were added to an automated sequential microwave digestion (CEM Mars 6 microwave digester). The resulting acidic solution was diluted to a volume of 50 mL using DI water (final concentration of the acid <5%). The ICP-OES was then calibrated with a series of matrix matched standards before introducing the unknown collagen samples. Digestion and ICP-OES analysis parameters are listed in **Supp. Tables 1 and 2** following similar protocols in literature [49, 60]. Six samples were used for each group and these were normalized to the calcium and phosphorous content of respective scaffolds post-hydration, crosslinking, and soaking without cells, to get a fold change and new calcium and phosphorous deposition. Additionally, an alkaline phosphatase (ALP) assay was used to determine hMSC cell-dependent mineralization in mineralized collagen scaffolds containing 0% honey, 5% incorporated honey, and 5% soaked honey. Phenol-red free media surrounding scaffolds for 14 days of hMSC cell culture on scaffolds was pooled and 80 μL of this sample was used with an Alkaline Phosphatase Assay Kit (Abcam, United Kingdom) along with phenol-red free media as a background control (n=6). ALP activity (U/mol) was calculated by the μmol/well of p-nitrophenylphosphate, reaction time, and sample volume.

### 2.10 Osteoprotegerin release from hMSC-seeded scaffolds

The amount of osteoprotegerin (OPG) released from hMSC-seeded scaffolds was quantified by an OPG ELISA (DY805, R&D Systems, Minnesota, USA). Media was collected every 3 days for 14 days and was pooled and assayed at specific timepoints: Day 3, Day 6 and 9, Day 12 and 14. 25 μL of sample was used and a media control was used to remove any background influence (n=6).

### 2.11 Endothelial cell tube formation assay

Human umbilical vein endothelial cells (HUVECs) (Lonza) were cultured in Endothelial Growth Medium 2 (EGM2, Lonza). Cells were maintained at 37 °C and 5% CO_2_ and were used before passage 5. To perform tube formation assay, 100 μL of growth factor reduced Matrigel (Corning, Tewksbury, MA) was dispensed per well in a 96-well plate and allowed to gel at 37 °C for one hour. 10,000 HUVECs were then mixed with 150 μL of each media sample per well (0, 5, 10, 25, 50 v/v% manuka honey dissolved in basal mesenchymal stem cell media) and pipetted over the Matrigel layer in each well (n=3). An additional experiment was performed with 10,000 HUVECs mixed with 150 μL of conditioned media generated by hMSCs cultured on mineralized collagen scaffolds with 0% honey, 5% incorporated honey, and 5% soaked honey (hMSC culture period of 6 days), with results compared to non-conditioned hMSC media as a control group (n=4). Tube formation was visualized and brightfield images were recorded using a DMi8 Yokogawa W1 spinning disk confocal microscope outfitted with a Hamamatsu EM-CCD digital camera (Leica Microsystems, Buffalo Grove, IL) or Leica DMI4000 B (Leica) at 6 and 12 hours. One to two regions were imaged per well.

### 2.12 Pseudomonas aeruginosa culture

*P. aeruginosa* PA14 [61] and PAO1 [62] were used throughout this study to quantify bacterial attachment and biofilm formation on mineralized collagen scaffolds. *P. aeruginosa* was grown at 37°C in lysogeny broth (LB, Difco LB Broth, Lennox Cat. No. 240230), Mueller-Hinton agar (MHA, Fisher Scientific) and KA medium, a modification of K10-T medium (no glycerol or tryptone, supplemented with 0.4% (wt/vol) L-arginine HCl) [63].

### 2.13 Zone of inhibition assay

Overnight cultures of *P. aeruginosa* strains PAO1 and PA14 were subcultured 1:100 in 2 mL LB and grown at 37°C until an OD_600_ of 1 was reached. Cultures were then normalized to OD_600_ of 0.5 and were streaked onto agar plates using a cotton swab and allowed to dry. Dry filter disks (6 mm diameter) soaked with 50 μL of 0, 5, 10, 25, and 50 v/v% manuka honey and scaffolds containing 0% honey, 5% incorporated honey, and 5% soaked honey were placed on the surface of the plates and incubated at 37°C for 16 hours.

### 2.14 Quantification of planktonic CFUs surrounding scaffolds

Overnight cultures of *P. aeruginosa* strains PAO1 and PA14 were subcultured 1:100 into 2 mL LB and grown at 37°C until an OD_600_ of 1 was reached. 100 μL of this culture was then centrifuged and the supernatant removed. The cell pellet was then resuspended in 10 mL KA medium resulting in a final OD_600_ of 0.01. 250 μL of OD_600_ 0.01 normalized culture was then added to each well of a 48-well plate containing a scaffold, resulting in a final inoculum of ~2.5×10^5^ CFU/well in 250 μL KA medium. Plates were then incubated, static and covered in a humidified chamber for 16 hours at 37°C. Following this incubation, serial dilutions of the bacteria and medium surrounding the scaffold were plated, incubated for 16 hours at 37°C, and counted to determine planktonic CFUs.

### 2.15 Assessment of gentamicin minimum bactericidal concentration

For our purposes, we defined minimum bactericidal concentration (MBC) as the lowest concentration of antibiotic in a 2-fold dilution series that resulted in no recovered planktonic CFU from growth medium after 6 hours of incubation. The initial inoculum was around 5 × 10^6 CFU / ml and the limit of detection of our CFU counts was 3.33 × 10^2 CFU / ml. The extent of killing with exposure to MBC gentamicin for 6 hours is therefore at least 99.99%.

Overnight cultures of *P. aeruginosa* strains PAO1 and PA14 were subcultured 1:100 into 2 mL LB and grown at 37°C until an OD_600_ of 1 was reached. 200 μL of this culture was then centrifuged and the supernatant removed. The cell pellet was then resuspended in 10 mL LB or KA medium resulting in a final OD_600_ of 0.02. 2-fold dilutions of gentamicin sulfate (Gold Bio, cat: G-400-5) ranging in concentration from 0.005 μg/ml to 102.4 μg/ml were prepared in either LB or KA medium and 100μl of was added to the wells of a 96-well plate. 100 μl of bacterial suspension was then added to each well. The plates were then covered and incubated static at 37°C in a humidified chamber for 6 hours. Planktonic cells were then enumerated by serially diluting and plating on LB plates followed by overnight incubation at 37°C and counting of colonies. The lowest concentration at which no colonies were observed was considered to be the MBC under these conditions.

### 2.16 Assessment of scaffold protection of *P. aeruginosa* from gentamicin

Overnight cultures of *P. aeruginosa* strains PAO1 and PA14 were subcultured 1:100 into 2 mL LB and grown at 37°C until an OD_600_ of 1 was reached. 100 μL of this culture was then centrifuged and the supernatant removed. The cell pellet was then resuspended in 10 mL LB or KA medium resulting in a final OD_600_ of 0.01. Scaffolds with either no honey, 5% honey incorporated into the scaffold, or that had been soaked in a 5% manuka honey solution as described in section 2.1, were added to wells of a 48-well plate. Immediately before adding bacteria to wells containing scaffolds, gentamicin stock solution was added to be bacterial suspension to produce the desired concentration. Three concentrations of antibiotic were used: no antibiotic, MBC, and 10X MBC (KA MBC: 0.4 μg/ml, KA 10X MBC: 4 μg/ml, LB MBC: 25.6 μg/ml, LB 10X MBC: 256 μg/ml). The bacterial suspension with antibiotic was then inverted several times and briefly vortexed to mix before 250 μl was added to each scaffold-containing well. The plates were then covered and incubated static at 37°C in a humidified chamber for 6 hours. Planktonic cells were then enumerated by serially diluting and plating on LB plates followed by overnight incubation at 37°C and counting of colonies.

### 2.17 Scanning Electron Microscopy of scaffold surface before and after bacterial culture

An Environmental Scanning Electron Microscope (FEI Quanta FEG 450 ESEM, FEI, Hillsboro, OR) was used to visualize the surface of scaffolds. Dry mineralized collagen scaffolds and 2% and 5% honey incorporated scaffolds prior to sterilization and crosslinking were cut in half to expose the inner structure of the scaffolds, and then were sputter coated for 70 seconds with Au/Pd using a Desk II TSC turbo-pumped sputter coater (Denton Vacuum, New Jersey, USA) before adding to the SEM. This was used to determine any structural and topographical changes in scaffold due to honey incorporating into the suspension. SEM images were also taken of scaffolds after culture in *P. aeruginosa*, including 0% honey, 5% honey incorporated, and 5% honey soaked scaffolds. Scaffolds were inoculated with *P. aeruginosa* strains PAO1 and PA14 following the method outlined in **Section 2.14** and **Section 2.16**. Following incubation, scaffolds were rinsed in sterile medium before adding to methacarn (methanol, chloroform, glacial acetic acid) for 2 hours at room temperature (**Section 2.14**) or methanol-free 4% formaldehyde (ThermoFisher, **Section 2.16**) and then kept in PBS at 4°C before washing in gradations of ethanol up to 100%. An Autosamdri 931 Critical Point Dryer (Tousimis, Maryland, USA) was used to dry samples to preserve bacterial structure and then dry scaffolds were sputter coated with Au/Pd for 70 seconds. To quantify bacteria numbers attached to the surface of the scaffold, 4 images were taken of the inside and outside of n=2 scaffolds and averaged.

### 2.18 Statistics

Statistical analysis of samples followed guidelines by *Ott and Longnecker* and previous experiments with similar sample groups [49, 57, 60, 64–66]. All samples were assessed for normality with a Shapiro-Wilk test and equal variance with a Levene test prior to statistical analysis. For comparisons of 3 or more samples, an ANOVA was used with a Tukey post-hoc for all data that had equal variance and normal distribution. Non-parametric ANOVAs were used for data that did not have equal variance or meet normality assumptions. For sample comparisons between 2 groups T-tests were used.

## 3. Results

### 3.1 Incorporation of manuka honey influences both mechanical properties and micoarchitecture of mineralized collagen scaffolds

The microstructural (porosity, SEM imaging) and mechanical (compression testing) properties of mineralized collagen scaffolds were characterized for conventional mineralized collagen scaffolds as well as scaffold variants containing 2% or 5% manuka honey (**Fig. 2**). SEM imaging demonstrated that as honey concentration was increased in mineralized collagen scaffolds, scaffolds exhibited a smoother surface; however, brushite mineral crystals were still visible within all scaffold variants. Scaffold porosity decreased with increasing amounts of incorporated honey, and interestingly the 2% manuka honey incorporated scaffolds displayed significantly increased Young’s modulus relative to the 0% and 5% Manuka honey scaffolds (**Fig. 2, Table 1**).

**Fig. 2.**
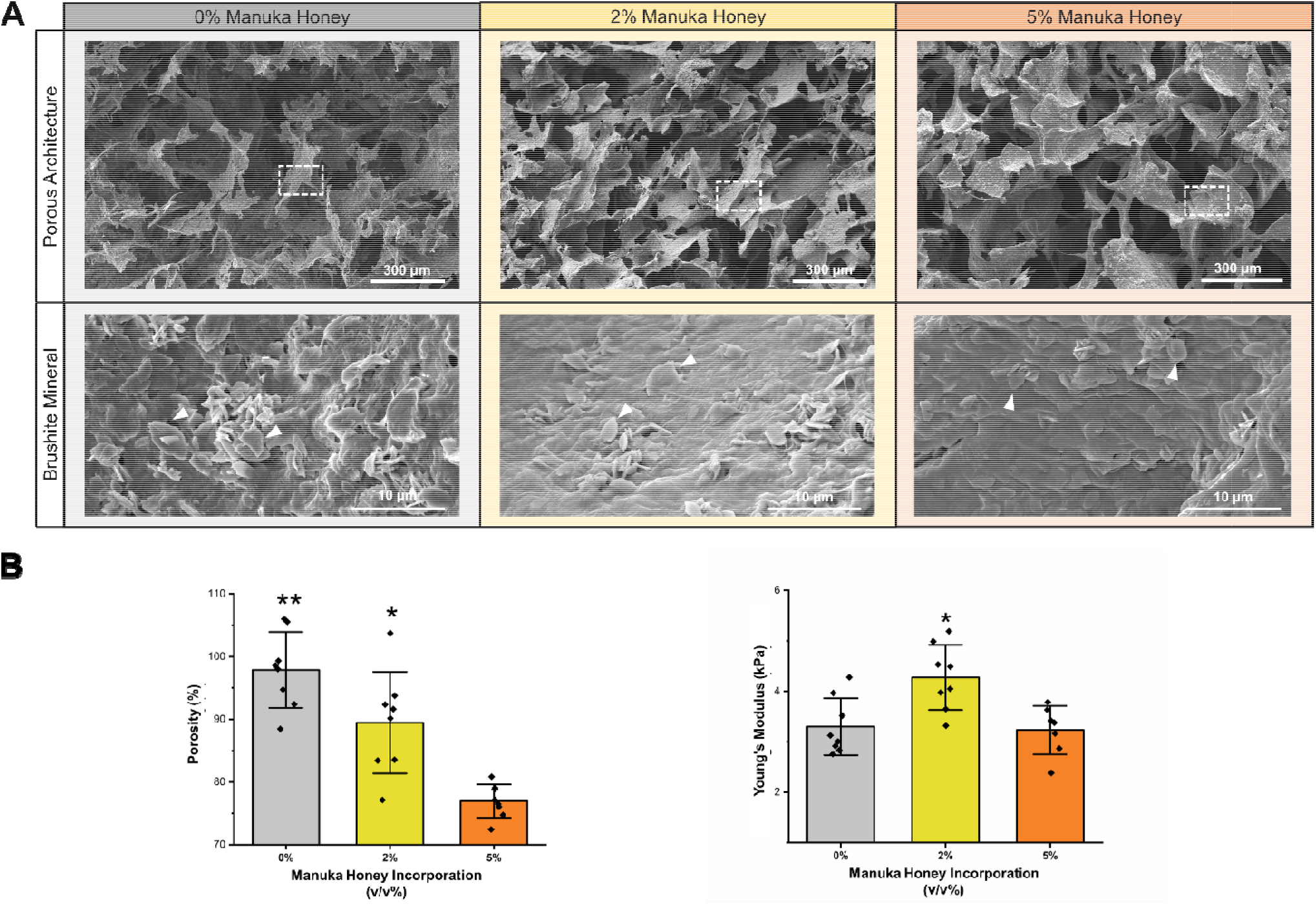
SEM images, porosity, and stiffness of mineralized collagen scaffolds incorporated with 0 v/v%, 2 v/v%, and 5 v/v% manuka honey. (A) SEM images of the internal structure of mineralized collagen scaffolds incorporated with manuka honey. The top row displays the porous architecture of each of the scaffold variants, also demonstrating that as more honey is added, the surface becomes smoother. The bottom row demonstrates that the brushite mineral crystals can be viewed in each of the scaffold variants as flat crystals and indicated by white arrowheads. (B) Porosity and Young’s Modulus of mineralized collagen scaffolds incorporated with manuka honey. Asterix designates significance, with all scaffold variants having significantly (p < 0.05) different porosities and the 2% manuka honey scaffolds having significantly different (p < 0.05) stiffness compared to the other two groups. Data expressed as average ± standard deviation (n=8).

**Table 1.**
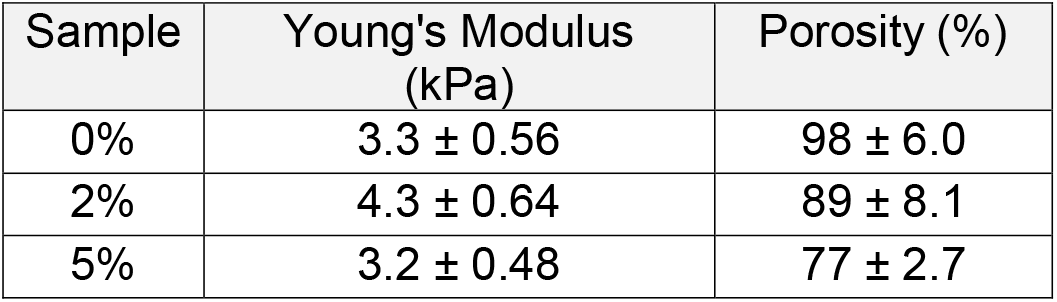
Young’s modulus and porosity of 0, 2, and 5 v/v% honey incorporated mineralized collagen scaffolds. Data represented as average ± standard deviation (n=8).

| Sample | Young's Modulus (kPa) | Porosity (%) |
| --- | --- | --- |
| 0% | 3.3 ± 0.56 | 98 ± 6.0 |
| 2% | 4.3 ± 0.64 | 89 ± 8.1 |
| 5% | 3.2 ± 0.48 | 77 ± 2.7 |

### 3.2 Honey soaked, but not honey incorporated, scaffolds enable glucose and MGO release

The amount of honey released from scaffolds incorporated and soaked with manuka honey was assessed via glucose and MGO release as well as via changes in pH. Scaffolds incorporated with honey during fabrication showed no appreciable burst release of glucose or MGO compared to conventional scaffolds without honey (**Fig. 3, Supp. Fig. 1**). There was a significantly greater amount of glucose and MGO released from the 5% honey soaked scaffolds compared to the mineralized collagen scaffolds without honey, however, the amount released was 13.6 times smaller than the MGO content in the 5 v/v% stock solution. Soaking scaffolds in 2% or 5% manuka honey did not significantly change the pH of the surrounding solution (**Supp. Fig. 1**).

**Fig. 3.**
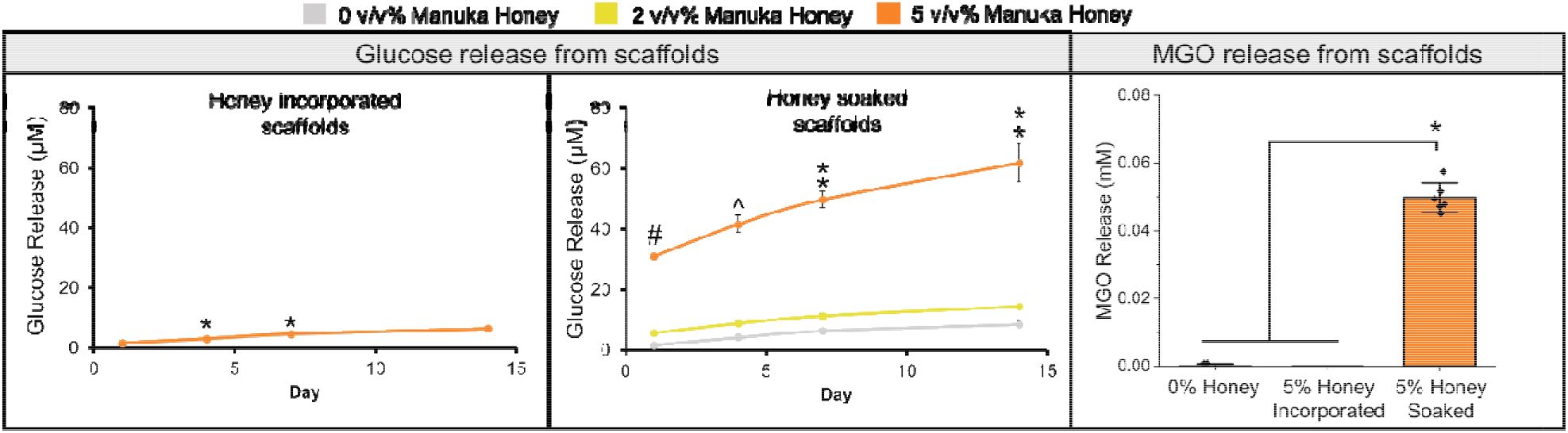
Glucose and methylglyoxal (MGO) release in mineralized collagen scaffolds incorporated and soaked in manuka honey. Glucose release was performed on samples over the course of 14 days soaked and incorporated in 2 and 5 v/v% manuka honey. MGO release was performed on 5 v/v% incorporated and 5 v/v% soaked samples after 1 day of soaking in PBS. The MGO concentration of the 5 v/v% manuka honey in PBS soaking stock solution was 0.68 mM. Asterix (*) designates significant (p < 0.05) differences in groups on the same day. Double asterix (**) indicates the 5 v/v% manuka honey group is significantly (p < 0.05) different from the other groups. Caret (^) indicates the 5 v/v% manuka honey group is significantly (p < 0.05) different from the 0 v/v% group. Hashtag (#) indicates all groups are significantly (p < 0.05) different from each other at the same timepoint. All data expressed as average ± standard deviation (n=6).

### 3.3 Incorporating large volumes (>10v/v%) of honey negatively affect hMSC metabolic health

Mineralized collagen scaffolds with 2 and 5% honey incorporated and 2, 5, 10, 25, and 50% honey soaked were assessed for hMSCs cell metabolic activity, or health, across 14 days. From day 3 onward, all cells seeded on 10, 25, and 50% honey soaked scaffolds had measured metabolic activities below 0, suggesting significant negative metabolic consequences of high levels of honey on cell health (**Fig. 4A**). After 7 days of hMSC culture on scaffolds, the 2% and 5% honey incorporated scaffolds had significantly (p < 0.05) reduced cell activity compared to 0% honey scaffolds (**Fig. 4B**). At days 1, 4, and 14, scaffolds soaked in 5% honey experienced significantly (p < 0.05) reduced cell activity compared to 2% and 0% honey soaked scaffolds (**Fig. 4C**). However, while lower than 0% (control) scaffolds 2% and 5% honey incorporated scaffolds still supported increased metabolic activity relative to initial seeding conditions.

**Fig. 4.**
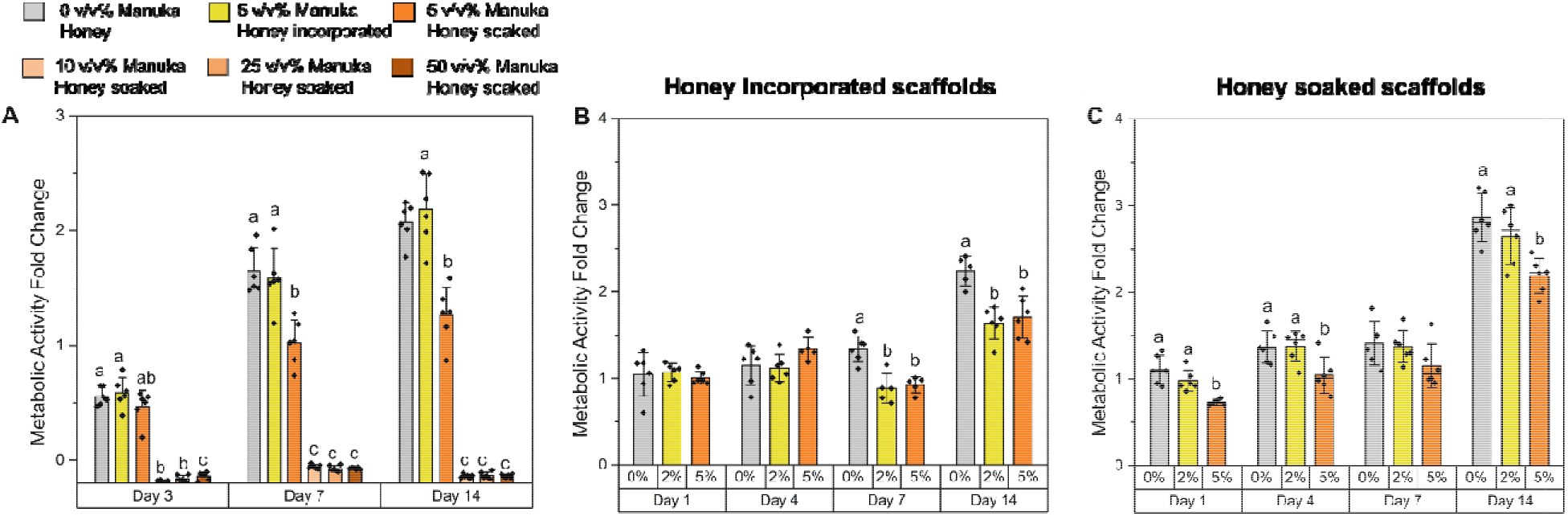
Metabolic activity of hMSCs on mineralized collagen-manuka honey scaffolds seeded with mesenchymal stem cells for 14 days. Alamar blue was used to assess the metabolic activity of 100,000 hMSCs seeded on scaffolds over the course of 14 days. A value of 1 represents the activity of 100,000 hMSCs before seeding on scaffolds. A value below 0 represents complete cell death. (A) Metabolic activity of hMSCs on 5 v/v% honey incorporated scaffolds and 0, 5, 10, 25, and 50 v/v% honey soaked scaffolds. (B) Metabolic activity of hMSCs on 0, 2, and 5 v/v% honey incorporated scaffolds. (C) Metabolic activity of hMSCs on 0, 2, and 5 v/v% honey soaked scaffolds. Different letters indicate significant (p < 0.05) differences between groups. All data expressed as average ± standard deviation (n=6).

### 3.4 Scaffolds soaked in 5% honey demonstrated greater mineral formation, alkaline phosphatase activity, and osteoprotegerin production

The osteogenic activity of hMSCs was assessed in honey containing scaffolds via calcium and phosphorous production, alkaline phosphatase content, and OPG release. 5% honey incorporated scaffolds demonstrated significantly (p < 0.05) reduced calcium and phosphorous content compared to 0 and 2% honey incorporated scaffolds after 7 and 14 days of culture (**Fig. 5**). 5% honey soaked scaffolds had the opposite trend, with a significant (p < 0.05) increase in calcium at day 14 and an increase in phosphorous at days 7 and 14 compared to conventional mineralized (0% honey) scaffolds. There were no differences in OPG production with the incorporation of honey into scaffolds, through 5% honey soaked scaffolds exhibited significant (p < 0.05) increase in OPG secretion by hMSCs at days 9 and 14 compared to 0 and 2% groups (**Fig. 5**). Both 5% honey incorporated and 5% honey soaked scaffolds had significantly (p < 0.05) higher alkaline phosphatase activity after 14 days of hMSC cell culture than the scaffolds without honey (**Fig. 5**).

**Fig. 5.**
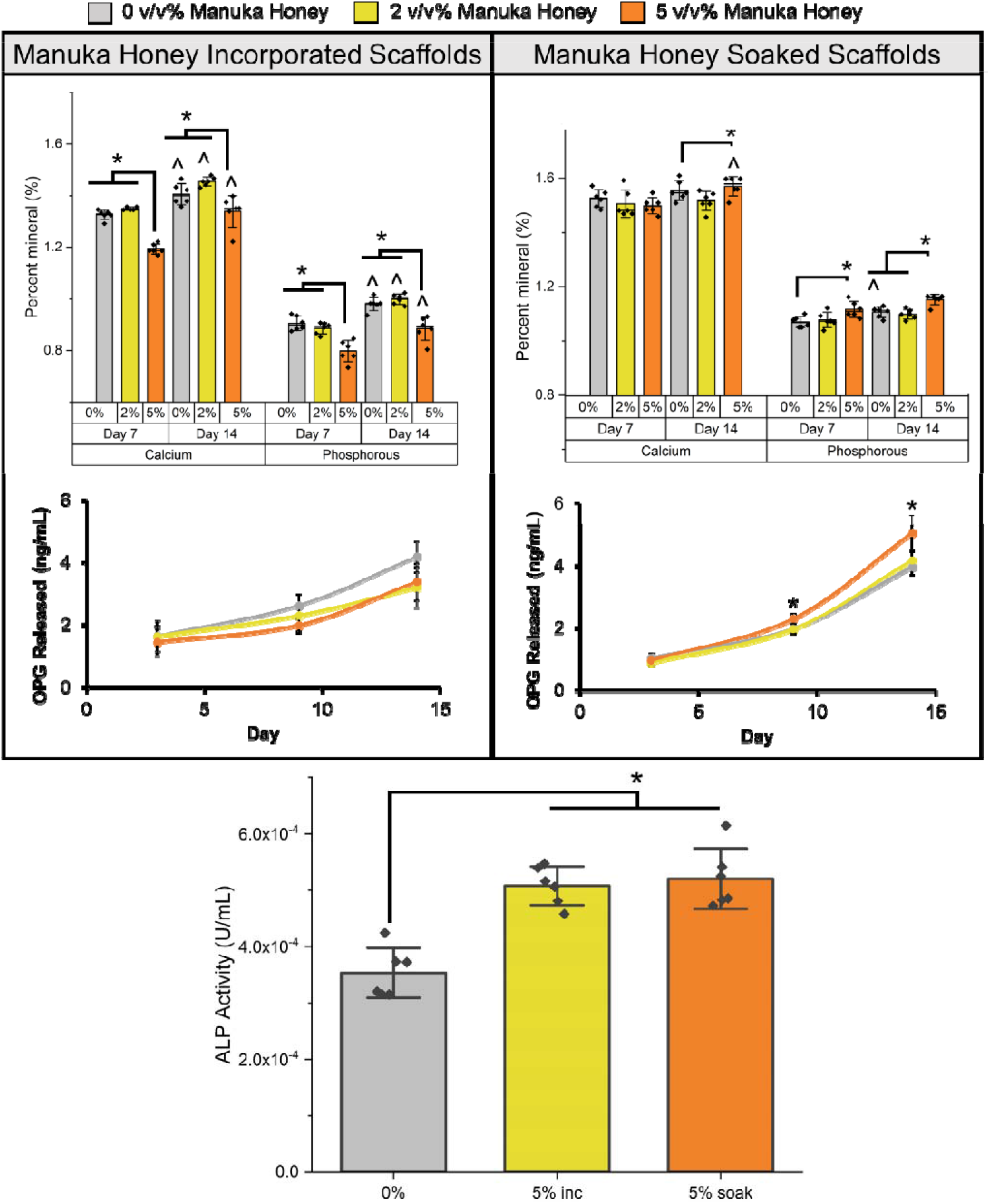
Mineral formation, osteoprotegerin release, and alkaline phosphatase (ALP) activity over 14 days of hMSC *in vitro* culture on mineralized collagen scaffolds incorporated and soaked with 2 and 5 v/v% manuka honey. For calcium and phospohorous percent data and ALP activity data, an asterix (*) denotes significant differences between indicated groups (p < 0.05) and a caret (^) designates significantly (p < 0.05) different values on day 14 compared to day 7. An asterix (*) on OPG release curves represents the 5 v/v% manuka honey group is significantly (p < 0.05) different from all other groups. All data expressed as average ± standard deviation (n=6).

### 3.5 While high levels of honey directly inhibit endothelial cell tube formation, MSC conditioned media generated in honey-functionalized scaffold does not

A Matrigel tube formation assay using embedded HUVECs was used to directly assess the consequence of honey (5, 10, 25, 50 v/v%) in basal mesenchymal stem cell media. Direct culture with manuka honey in the media inhibited vessel formation (**Supp. Fig. 2**), as evidenced by a lack of cell networks in media containing concentrations of 5% honey and above. However, release profiles of honey incorporated or honey soaked scaffolds suggested significantly lower amounts of honey were released into the media (**Fig. 3**). As a result, endothelial network formation was assessed in response to conditioned media generated by hMSCs cultured in mineralized collagen scaffolds (0%; 5% incorporated; 5% soaked) for 6 days. Endothelial networks were visible in all groups at 6 hours (**Fig. 6A, Supp. Fig. 3**), with no significant (p < 0.05) differences in the ability of HUVECs to form tubes in all groups compared to unconditioned media controls (**Fig. 6B**).

**Fig. 6.**
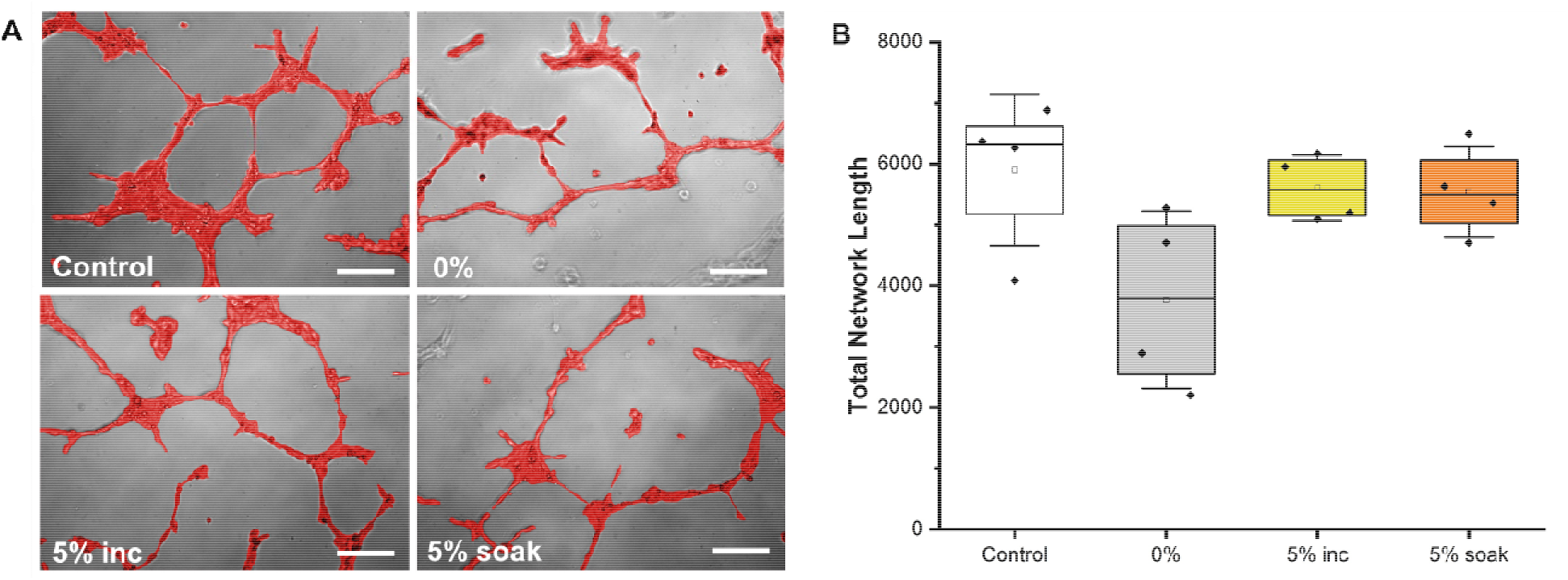
Human umbilical vein endothelial cell (HUVEC) tube formation assay on Matrigel after 6 hours with conditioned media from mineralized collagen scaffolds seeded with hMSCs (6 days) with 0% manuka honey, 5% manuka honey incorporated, 5% manuka honey soaked, and hMSC media without conditioning as a control (0%, 5% inc, 5% soak, Control, respectively). (A) Brightfield images of HUVECs forming tubes, false colored red, scale bar represents 200 μm. Original image can be viewed in **Supp. Fig. 3**. (B) Box plot representing the total network length from ImageJ analysis of brightfield images of HUVEC tubes. No significance (p < 0.05) between groups. Data represented as median (line), mean (white dots), individual data points (black dots), min and max (whiskers).

### 3.6 P Honey soaked and incorporated scaffolds and high concentrations of honey soaked filter discs do not inhibit P. aeruginosa growth

Filter discs soaked in 5, 10, 25, and 50 v/v% manuka honey were added to lawns of *P. aeruginosa* and the zone of inhibition from these was assessed. All concentrations of honey demonstrated no zones of inhibition, even concentrations as high as 50 v/v% (**Supp. Fig. 4**). Similarly, mineralized collagen scaffolds with 0% honey, 5% honey incorporated, and 5% honey soaked demonstrated no zones of inhibition (**Supp. Fig. 5**). These data suggest that *P. aeruginosa* adjacent to the scaffold are not sensitive to the release of manuka honey, but motivated studies to evaluate growth and biofilm formation within the scaffold.

### 3.7 Incorporating honey into scaffolds does not prevent bacterial attachment and soaking scaffolds in honey prevents attachment but increases bacterial growth in the surrounding medium

SEM images of the surface of scaffolds cultured with PA14 and PAO1 *P. aeruginosa* after 6 and 16 hours demonstrated bacterial attachment regardless of honey soaking or incorporation (**Fig. 7A, Supp. Fig. 6-8**). While the 0% and 5% honey incorporated groups had little to no free scaffold visible and were coated in bacteria, 5% honey soaked scaffolds displayed areas of visible mineralized collagen scaffold, suggesting honey soaking may inhibit bacteria attachment to the scaffold surface (**Fig. 7A**). Quantification of the CFUs of planktonic bacteria in the medium surrounding the scaffolds after 16 hours of incubation revealed that 5% honey soaked scaffolds had significantly (p < 0.05, PA14; p < 0.001, PAO1) greater planktonic CFUs (**Fig. 7B**). There were no differences in planktonic CFUs between the 0% (control) and 5% incorporated honey scaffolds in the surrounding medium.

**Fig. 7.**
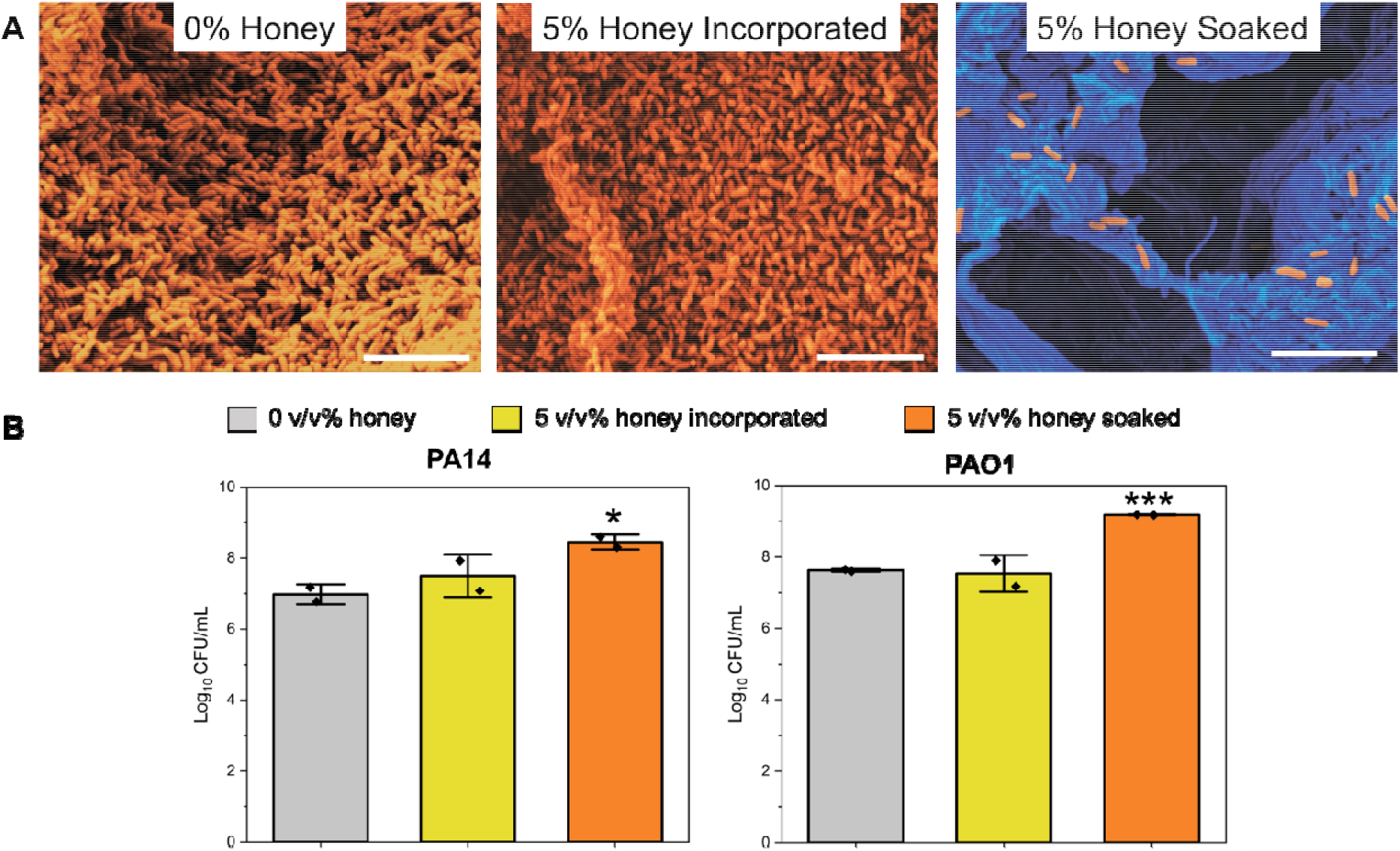
Assessment of *P. aeruginosa* growth and attachment on mineralized collagen scaffolds with manuka honey. Scaffolds containing 0% honey, 5% incorporated honey, and 5% soaked honey were cultured in medium containing PA14 or PAO1 *P. aeruginosa* strains for 16 hours. (A) SEM images demonstrating attachment of *P. aeruginosa* on scaffolds after 16 hours of PA14 culture. Images are false colored to demonstrate bacteria presence on scaffold surface. Orange represents *P. aeruginosa* bacteria and blue represents mineralized collagen scaffold. Scale bar represents 5 μm. Original SEM images can be viewed in **Supp. Fig. 6-8**. (B) Quantification of *P. aeruginosa* growth in medium containing scaffolds with or without honey treatment. 48-well plates containing scaffolds were inoculated with *P. aeruginosa* PAO1 and PA14 and incubated statically for 16 hours. Planktonic populations were recovered from the medium surrounding the scaffolds and CFU were counted. Planktonic CFU are presented per well. Each column represents the average of two biological replicates, each performed using three technical replicates. Error bars represent the standard deviation. Asterisks indicate values that differ significantly from CFU counts from the 0% honey scaffold treatment. *, P < 0.05, ***, P < 0.001, two-tailed student’s t-test. Data represented as average ± standard deviation.

### 3.8 P. aeruginosa is equally sensitive to killing by gentamicin regardless of honey treatment of scaffolds

Next, we investigated the sensitivity of *P. aeruginosa* to gentamicin when incubated with a scaffold with or without manuka honey. The planktonic MBC for gentamicin was found to be 0.4 μg/ml in KA medium, and 25.6 μg/ml in LB medium. We inoculated bacteria with either no gentamicin, the MBC of gentamicin, or 10 times the MBC of gentamicin onto scaffolds with no honey, 5% honey incorporated scaffolds, or 5% honey soaked scaffolds. We assessed survival in the medium using planktonic CFUs, and bacterial attachment and survival on the scaffold surface using SEM (**Fig. 8**).

**Fig. 8.**
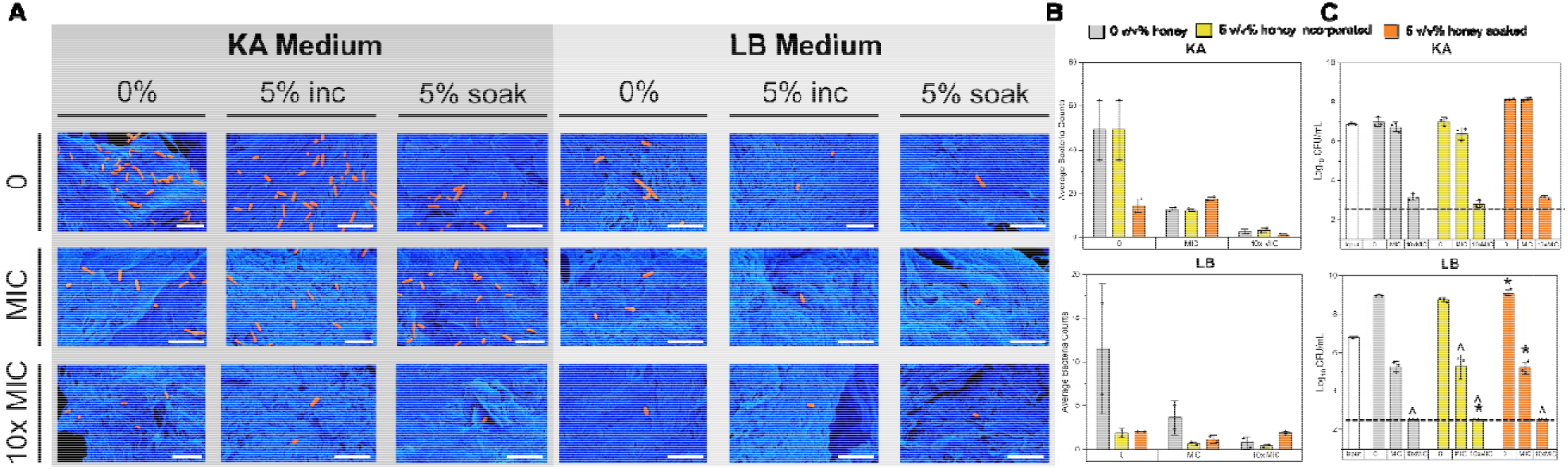
Quantification of bacterial attachment and planktonic activity after 6 hours of PA14 *P. aeruginosa* culture on mineralized collagen scaffolds with 0% honey (0%), 5% incorporated honey (5% inc), and 5% soaked honey (5% soak) with no additional antibiotics (0), minimum inhibitory concentration (MBC) with gentamicin, and 10 times the MBC (10x MBC). Scaffolds were cultured in KA medium and lysogeny broth (LB) medium. (A) SEM images were taken of the surface of scaffolds, false-colored to better visualize bacteria (orange) against the scaffold (blue). Scale bar represents 5 μm, uncolored images can be found in **Supp. Fig. 9**. (B) Attached bacteria in SEM images were counted and averaged across groups. No significant difference was found between groups (0%, 5% inc, 5% soak, n=2). Data expressed as average ± standard deviation. (C) Planktonic CFU counts of bacteria solution added to scaffolds (input) and measurements after 6 hours of incubation. The dashed line represents the limit of detection of the assay. For the LB graph, all groups are significantly (p < 0.05) different from the input, and between the 0, MIC, 10x MIC for one group (0%, 5% inc, 5% soak), but no significance was found between the 0%, 5% inc, and 5% soak groups between the different gentamicin concentrations. For the KA graph, * indicates significant (p < 0.05) differences between the indicated group and all other groups (including input) at the indicated gentamicin concentration. ^ indicates the group is significantly (p < 0.05) different from the 0 group for the same sample group (0%, 5% inc, 5% soak). Data expressed as average ± standard deviation (n=3 biological replicates).

We observed no difference in the number of CFUs recovered when incubated in KA medium with either no gentamicin or the MBC of gentamicin. However, when cells were grown in KA with 10X MBC of gentamicin, there were significantly fewer CFUs recovered. Interestingly, the number of CFUs recovered was not significantly different between the plain scaffold, 5% honey incorporated scaffold, and 5% soaked scaffold. No significant differences were found between the scaffold groups in terms of attached bacteria numbers, and bacteria were still present even in the 10x MBC group. In LB medium, we recovered significantly fewer CFUs when cells were incubated with the MBC of gentamicin versus when they were incubated in the absence of antibiotic. In the presence of 10X MBC gentamicin, no CFUs were recovered from any scaffold condition in LB medium. SEM imaging demonstrated fewer attached bacteria in the antibiotic-free group when scaffolds were cultured in LB medium than when cultured in KA medium. Finally, while no planktonic CFUs were recovered when bacteria were incubated in LB medium with 10X MBC, bacteria were still present on the surface of the scaffold.

## 4. Discussion

Regeneration of CMF defects requires biomaterials with innate osteogenic and antimicrobial properties. Here, we examined the effectiveness of manuka honey in mineralized collagen scaffold to reduce *P. aeruginosa* attachment and the effect of incorporated honey on osteogenesis of hMSCs. We hypothesized that mineralized collagen scaffolds do not resist bacterial attachment and biofilm formation, and manuka honey could be added to prevent this attachment.

We first investigated two disparate approaches to incorporate manuka honey into mineralized collagen scaffolds: direct incorporation during fabrication or soaking after scaffold fabrication. Increasing honey incorporation into the scaffolds resulted in an altered scaffold topography, including a reduced porosity. While incorporation of 2% honey into scaffolds resulted in increased scaffold stiffness, the increase was only incremental. The amount of honey released from honey incorporated and honey-soaked scaffolds was assessed by glucose and MGO. 2 and 5% honey incorporated scaffolds had the same glucose and MGO release as 0% honey scaffolds, suggesting little to no honey release from these scaffolds. However, 2 and 5% honey soaked scaffolds displayed increased glucose release, with the 5% soaked group having the highest amount of glucose released and higher MGO released than the 0% and 5% honey incorporated scaffolds. Release of glucose and MGO suggest that soaking already fabricated scaffolds in honey solution was the better method to facilitate release of honey from the scaffold into the surroundings; however, quantifying the total amount of honey released also suggested that even for honey soaked scaffolds, the majority of the honey was retained in the scaffold. These findings were also consistent with prior work to sequester growth factors within the mineralized collagen scaffolds, which also identified a significant fraction was retained within the scaffold [57]. The pH of honey soaked and incorporated scaffolds was also assessed, as mineralized collagen scaffolds have been known to release mineral over time and causing a reduction in pH, with honey also known to have a low pH [30, 60]. Over the course of 14 days there were few relative changes in pH (average of all groups: 6.5 ± 0.0048) where neither honey soaking or incorporation leading to reduced pH. This could be beneficial, as acidic pH below 7 may lead to bone resorption [67, 68].

hMSC metabolic activity was measured in response to increasing concentrations of manuka honey soaked into scaffolds (5, 10, 25, 50%). After 3 days of cell culture, the metabolic activity of hMSCs seeded on 10, 25, and 50% honey soaked scaffolds was not measurable, suggesting cells may be either dead or metabolically suppressed. This correlates with previous studies, where a 5 v/v% or greater concentration of manuka honey on fibroblasts, pulmonary microvascular endothelial cells, and macrophages killed nearly all these cells after one day of culture [68, 69]. Additionally, 2% and 5% honey incorporated scaffolds had lower metabolic activity than scaffolds without honey at days 7 and 14, and 5% honey soaked scaffolds had consistently lower activity than scaffolds without honey. However, hMSCs in these scaffolds did show an increase in metabolic activity over time. More importantly, 5% honey soaked scaffolds displayed greater calcium and phosphorous mineral formation and alkaline phosphatase activity than other groups (including 0% honey control scaffolds). Increased alkaline phosphatase activity in 5% honey soaked variants suggests more cell-dependent active bone formation, which may explain the decrease in metabolic activity if these cells were producing mineral and differentiating as opposed to expanding. This is consistent with prior studies filling small mandibular bone defects in rats with honey, which found faster mineralization [70]. Decreased calcium and phosphorous production in the 5% honey incorporated scaffolds could be attributed to decreased porosity and decreased cell activity in this group. Soaking scaffolds in 5% honey also led to higher osteoprotegerin (OPG) release, a glycoprotein responsible for blocking the ability of osteoclast precursors to differentiate into osteoclasts and resorb bone. We previously showed mineralized collagen scaffolds endogenously promote OPG production by seeded osteroprogenitors [71, 72], so this finding that honey incorporation promotes even greater OPG production is significant. An increase in OPG could help to maintain the balance of osteoclasts and osteoblasts in bone homeostasis even during infection and apoptosis of some osteoblasts and would be worthy of future study to investigate MSC-osteoclast crosstalk in mineralized collagen scaffolds as a function of inflammatory challenge.

We subsequently observed two distinct phenotypes in regards to endothelial tube formation. Using a Matrigel assay, we visualized tube formation in media without any honey, but culture with 5, 10, 25, or 50% manuka honey in media completely inhibited tube formation. This inhibition could have been due to cell death, which as previously mentioned, concentrations 5 v/v% and above resulted in complete pulmonary microvasculature endothelial cell death [68, 69]. As we observed that mineralized collagen scaffolds soaked in 5% manuka honey did not release all 5% of this honey to the surrounding medium, it is likely endothelial cells in the scaffold microenvironment may experience lower concentrations. To test this, we performed an additional tube formation assay using conditioned media generated by hMSCs seeded on mineralized collagen scaffolds (0% honey, 5% incorporated honey, and 5% soaked honey) for 6 days. Here, 5% soaked or incorporated honey scaffolds did not negatively impact endothelial tube formation, with HUVECs assembling network structures in all cases. This suggests addition of honey to the scaffolds did not functionally reduce endothelial tube formation potential.

Designing scaffolds to resist bacteria attachment may further improve patient outcomes, as infection can significantly inhibit bone regeneration. Closed fractures without biomaterial intervention have much lower infection rates (1.5%), however, open fractures such as CMF defects requiring biomaterial intervention have infection rates ranging from 3-40%, and treatment of these will likely exceed $11 billion in the US annually with a growing population [6–8]. This represents a clinical problem in biomaterial design, as antibiotics may be able to kill non-adherent bacteria, but once a biofilm has formed on implant surfaces, antibiotics are less effective at clearing the infection due to the increased antibiotic tolerance of bacterial cells in a biofilm [16]. A previous study showed for both PA14 and PAO1 strains of *P. aeruginosa*, planktonic bacteria can be completely eradicated and biofilms reduced by a concentration of 16% honey in the growth medium [73]. Although the concentration of honey used in our study is lower than that found to be effective in this previous study, we sought to assess the antimicrobial potential of manuka honey in collagen scaffolds. We saw no zones of inhibition resulting from honey containing scaffolds and discs, indicating the concentration of honey released from 5% honey incorporated or soaked scaffolds is insufficient to reduce *P. aeruginosa* viability. The ability of honey to inhibit bacterial attachment was then assessed by visualizing attached cells on the surfaces of scaffolds soaked in KA medium containing 0% honey, 5% incorporated honey, or 5% honey soaked scaffolds. Only these concentrations were used as prior data indicate that higher concentrations negatively affected hMSCs, and lower concentrations were hypothesized to be less likely to alter bacterial colonization. After incubation of *P. aeruginosa* (PA14, PAO1) with scaffolds for 6 or 16 hours, SEM imaging of the scaffold surface demonstrated bacterial attachment on all scaffold types. In particular, the 0% honey and 5% honey incorporated scaffold surfaces were entirely covered in PA14 bacteria after 16 hours, supporting our hypothesis that mineralized collagen scaffolds have no native method to reduce bacterial attachment and would be prone to colonization by bacteria present in the wound site. Of significance, 5% honey soaked scaffolds had noticeably fewer bacteria adhered to the surface, as the mineralized collagen scaffold was still visible in images of the surface, but still substantial bacteria attached in PA14 and PAO1 strains. A previous study demonstrated that low concentrations of honey (< 2%) could increase biofilm formation by PAO1 [73]. The 5% honey <u>soaked</u> scaffolds were likely increasing the overall concentration of honey in the growth medium to < 2% honey. However, we did not observe an increase in biofilm formation as was reported in that study. Any ability of scaffolds to reduce bacterial attachment could represent a large impact on improving infection clearance, as this represents the significant issue in CMF infections requiring surgical intervention. Quantification of planktonic bacteria in the medium surrounding the scaffold after 16 hours revealed that bacterial growth was greater when incubated with 5% honey soaked scaffolds. Previous studies have highlighted the complex role of nutritional cues in biofilm formation and maintenance [74, 75]. The increase in planktonic CFUs and decrease in observed surface attachment visible after 16 hours of bacterial culture, are consistent with a model whereby honey may reduce surface attachment and biofilm formation by impacting *P. aeruginosa* metabolism.

The observation that soaking scaffolds in 5% honey results in increased bacterial growth in KA medium, suggests the potential of worse patient outcomes. However, prophylactic antibiotic treatment such as gentamicin commonly follows bone graft or biomaterial implantation [76, 77]. We therefore sought to determine whether soaking scaffolds in 5% honey would be likely to increase the bacterial burden even in the presence of antibiotics or, conversely, if honey may increase the sensitivity of *P. aeruginosa* to antibiotics. We found no evidence that 5% honey incorporated or 5% honey soaked scaffolds increased the sensitivity of *P. aeruginosa* to gentamicin (**Fig. 8**). Instead, we observed that gentamicin reduced *P. aeruginosa* viability to the same extent in all three scaffold conditions tested, suggesting the increased bacterial growth observed when incubated with the 5% honey soaked scaffold may not translate to an increased bacterial burden in a graft-site infection. Furthermore, the increase in bacterial growth when incubated with 5% honey soaked scaffolds was only seen in KA medium, which has very little carbon to support growth, but not in LB, which is a comparably rich medium. Whether honey would promote bacterial growth in a wound site is unclear. Additionally, adding gentamicin antibiotic to the 0% honey, 5% incorporated, and 5% soaked groups cultured in PA14 for 6 hours demonstrated a drop in attached bacteria, but not complete inhibition, even with the addition of 10x the MBC. This demonstrated that addition of manuka honey did not substantially prevent *P. aeruginosa* attachment, and scaffolds may protect bacteria from complete killing by antibiotics.

There were a few limitations to this bacterial study. Bacterial attachment on scaffolds was not quantitatively assessed due to the difficulty of recovering accurate cell counts from biofilms within complex three-dimensional surfaces. Further, *P. aeruginosa* is not commonly isolated from bone infections and so the impact of manuka honey on *P. aeruginosa* colonization of bone grafts is not as clinically relevant as the impact on a more commonly isolated organism, such as *S. aureus*. Future work aims to test these scaffolds with *S. aureus* bacteria including Methicillin-Resistant Staphylococcus aureus (MRSA). *S. aureus* and antibiotic resistant strains of *S. aureus* are the most common bacteria present in bone infection and *S. aureus* contributes to (or is isolated from/detected in) four out of five implant-associated infections [78, 79]. *S. aureus* not only causes abscess formation in soft-tissue infections, but also directly interacts with and infects osteoblasts. Infection of osteoblasts can cause apoptosis and inadvertently increase bone resorption through an imbalance in osteoclast and osteoblast activity [80]. Recently, *S. aureus* culture on human bone samples demonstrated a significant drop in bone mineral quality and crystallinity, as well as altered collagen cross-linking [81], and in *in vivo* studies, localizes to the canaliculi of infected bone [82, 83]. We expect 5% honey soaked scaffolds to prevent bacterial attachment of *S. aureus* as past work with manuka honey in broth cultures demonstrated 10.8 v/v% honey was needed for 100% growth inhibition of *P. aeruginosa*, while only 1.8 v/v% honey was needed to completely inhibit *S. aureus* cultures [22]. Additionally, we plan to study the impact of *S. aureus* on osteogenesis of mesenchymal stem cells and immune cells on mineralized collagen scaffolds, if honey can promote osteogenesis even in the presence of bacteria, and if honey can suppress a pro-inflammatory immune response [70]. Other scaffolds fabricated with manuka honey have demonstrated clearance of *S. aureus* at higher concentrations >10%, however, the effect of this concentration on the regeneration of bone and bone cell health has not been investigated [23, 26]. Additionally, there is significant opportunity to better understand the impact of honey on endothelial vasculature formation using a range of recent advances in three-dimensional vessel imaging [84] and quantitative assessment [85].

## 5. Conclusions

We investigated the potential for manuka honey in mineralized collagen scaffolds to inhibit *P. aeruginosa* attachment, mesenchymal stem cell osteogenesis, and angiogenic potential. While we developed both incorporated and soaked approaches to add manuka honey into mineralized collagen scaffolds, only honey soaked scaffolds released honey into the surroundings. Soaking concentrations of manuka honey 10% and above resulted in mesenchymal stem cell death. Although soaking scaffolds in 5% manuka honey decreased hMSC metabolic activity, more phosphorous and calcium mineral was produced, more osteoprotegerin was released, and these scaffolds had higher alkaline phosphatase activity over 14 days of culture than scaffolds without honey. Furthermore, incorporation and soaking of scaffolds in 5% manuka honey did not negatively impact the ability of endothelial cells to form tubes. Finally, addition of manuka honey did not have an impact on *P. aeruginosa* attachment but may have a potential for other bacteria strains. This work demonstrated that soaking mineralized collagen scaffolds in 5 v/v% manuka honey could increase the ability of these scaffolds to produce mineral and promote osteogenesis, while potentially reducing bacterial attachment and subsequent biofilm formation.

## Acknowledgements

I would like to acknowledge the following institutes for access to their facilities and services: the School of Chemical Sciences Microanalysis Laboratory, the Carl R. Woese Institute for Genomic Biology, and the Beckman Institute for Advanced Science and Technology, located at the University of Illinois. I would also like to acknowledge Scott Robinson and Cate Wallace for assistance with critical point drying and SEM imaging of samples. Research reported in this publication was supported by the National Institute of Dental and Craniofacial Research of the National Institutes of Health under Award Number R21 DE026582 and R01 DE030491 (BACH). We are also grateful for funds provided by the NSF Graduate Research Fellowship DGE-1144245 (MJD), the Allen Distinguished Investigator Award (AJC), and R01 AR076941 (NJH) to perform this research. The interpretations and conclusions presented are those of the authors and are not necessarily endorsed by the National Institutes of Health or the National Science Foundation.

## Contributions (CRediT: Contributor Roles Taxonomy [86, 87])

**Marley Dewey:** Conceptualization, Data curation, Formal Analysis, Visualization, Investigation, Methodology, Writing – original draft, Writing – review & editing. **Alan Collins:** Resources, Data curation, Methodology, Formal Analysis, Investigation, Writing – review & editing. **Victoria Barnhouse:** Investigation, Formal Analysis. **Aleczandria Tiffany:** Investigation, Visualization **Crisyln Lu:** Investigation. **Vasiliki Kolliopoulos:** Investigation.. **Noreen Hickok:** Conceptualization, Methodology, Writing – review & editing. **Brendan Harley:** Conceptualization, Resources, Project administration, Funding acquisition, Supervision, Writing – review & editing.

## Data Availability

The raw and processed data required to reproduce these findings will be available to download upon publication of this manuscript: Dewey, Marley (2022), “Evaluation of P. aeruginosa attachment on mineralized collagen scaffolds and addition of manuka honey to increase mesenchymal stem cell osteogenesis”, Mendeley Data, V1, doi: 10.17632/kx34gyyr2x.1.

