## Supplementary Files for "Evaluation of *P. aeruginosa* attachment on mineralized collagen scaffolds and addition of manuka honey to increase mesenchymal stem cell osteogenesis"

^1^ Dept. of Materials Science and Engineering

^2^ Carl R. Woese Institute for Genomic Biology

^3^ Dept. of Chemical and Biomolecular Engineering

^4^ Dept. of Bioengineering

^5^ School of Chemical Sciences

^7^ Cancer Center at Illinois

University of Illinois Urbana-Champaign

Urbana, IL 61801

^6^ Department of Orthopaedic Surgery

Sidney Kimmel Medical College of Thomas Jefferson University

Philadelphia, PA 19107

**Corresponding Author:**

B.A.C. Harley

Dept. of Chemical and Biomolecular Engineering

Carl R. Woese Institute for Genomic Biology

Cancer Center at Illinois

University of Illinois at Urbana-Champaign

110 Roger Adams Laboratory

600 S. Mathews Ave.

Urbana, IL 61801

**Supp. Table 1** Parameters for microwave assisted acid digestion of collagen samples (see **section 2.9**).

| Digestion Parameters | Values |
| --- | --- |
| Power | 1800 W |
| Temperature | 200 ^0^C |
| Ramp Time | 35 minutes |
| Hold Time | 25 minutes |

**Supp. Table 2** Parameters for analysis of calcium and phosphorus collagen samples via Optima 8300 ICP-OES (see **section 2.9**). Emission lines: Ca ((II)-317.93 nm) and P ((I)- 213.62 nm).

| ICP-OES Parameters | Values |
| --- | --- |
| RF Power | 1500 Watts |
| Nebulizer | GemCone Low Flow |
| Nebulizer Gas Flow rate | 0.85L/min |
| Plasma Gas Flow rate- Argon | 10L/min |
| Sample Flow rate | 1.50mL/min |

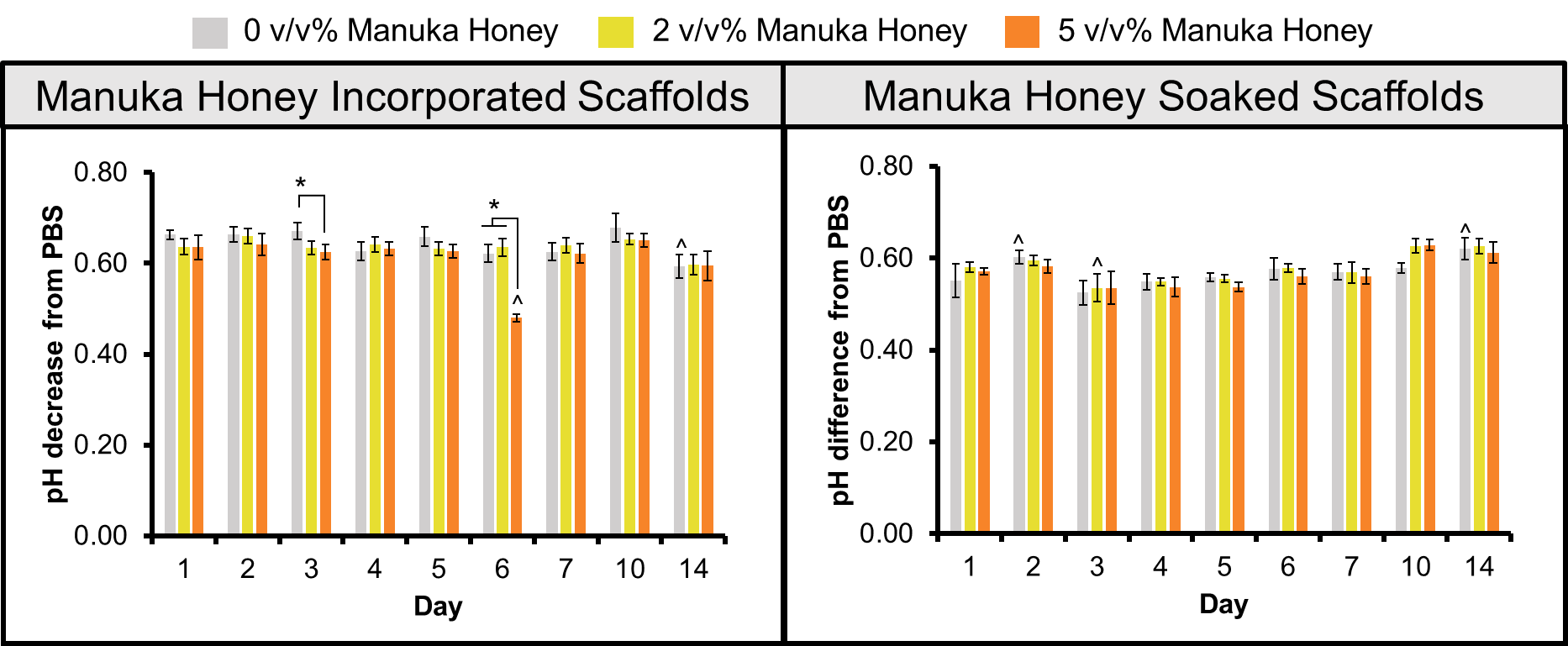

**Supp. Fig. 1** pH changes of 2 and 5 v/v% manuka honey soaked and incorporated scaffolds over 14 days. Scaffolds were soaked in PBS and the pH was measured at days 1-7, 10, and 14. The y-axis represents the pH drop at each of the timepoints (PBS pH averaged 7.12). Asterix (*) represents significant (p < 0.05) differences between indicated groups. Carrot (^) represents significant (p < 0.05) change in pH of one group compared to the same group on day 1. All data expressed as average ± standard deviation (n=6).

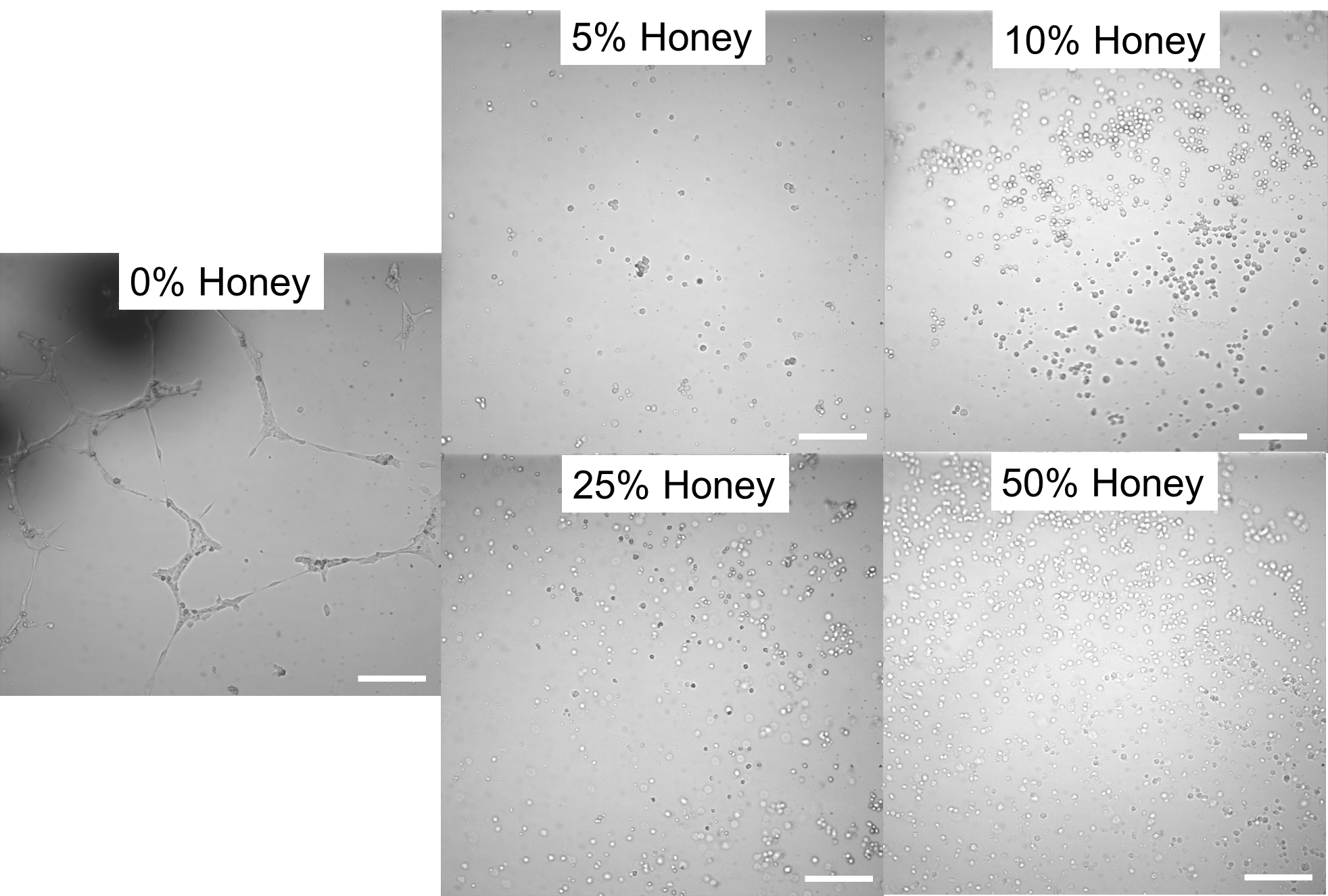

**Supp. Fig. 2** Images of human umbilical vein endothelial cell tube formation after 6 hours of culture in a Matrigel assay with 0%, 5%, 10%, 25%, and 50% honey in complete mesenchymal stem cell media. Scale bar represents 200 µm (n=3).

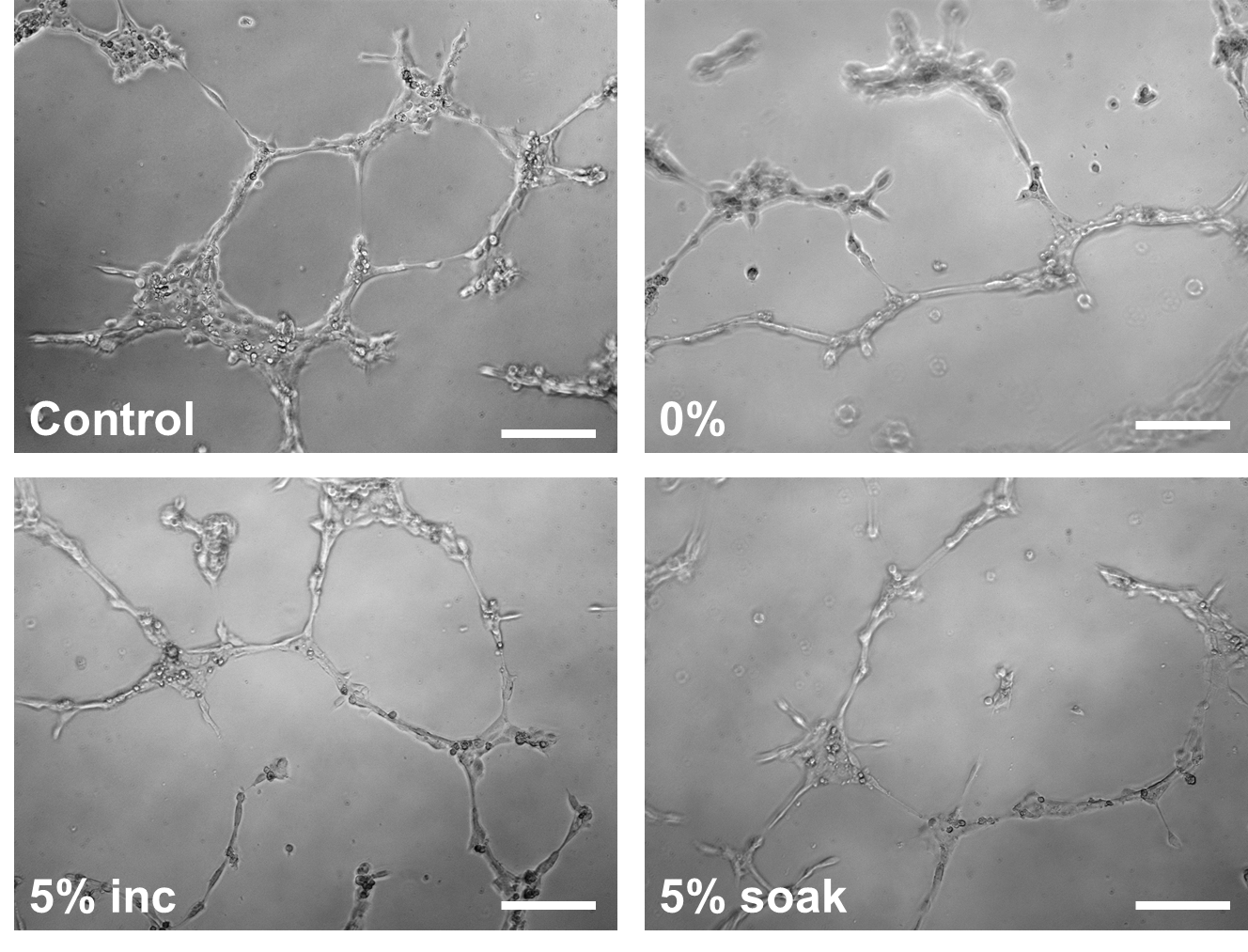

**Supp. Fig. 3.** Endothelial cell tube formation assay on Matrigel after 6 hours with conditioned media from mineralized collagen scaffolds seeded with hMSCs with 0% manuka honey, 5% manuka honey incorporated, 5% manuka honey soaked, and hMSC media without conditioning as a control (0%, 5% inc, 5% soak, Control, respectively). Brightfield images of HUVECs forming tubes, scale bar represents 200 µm, see Figure 6.

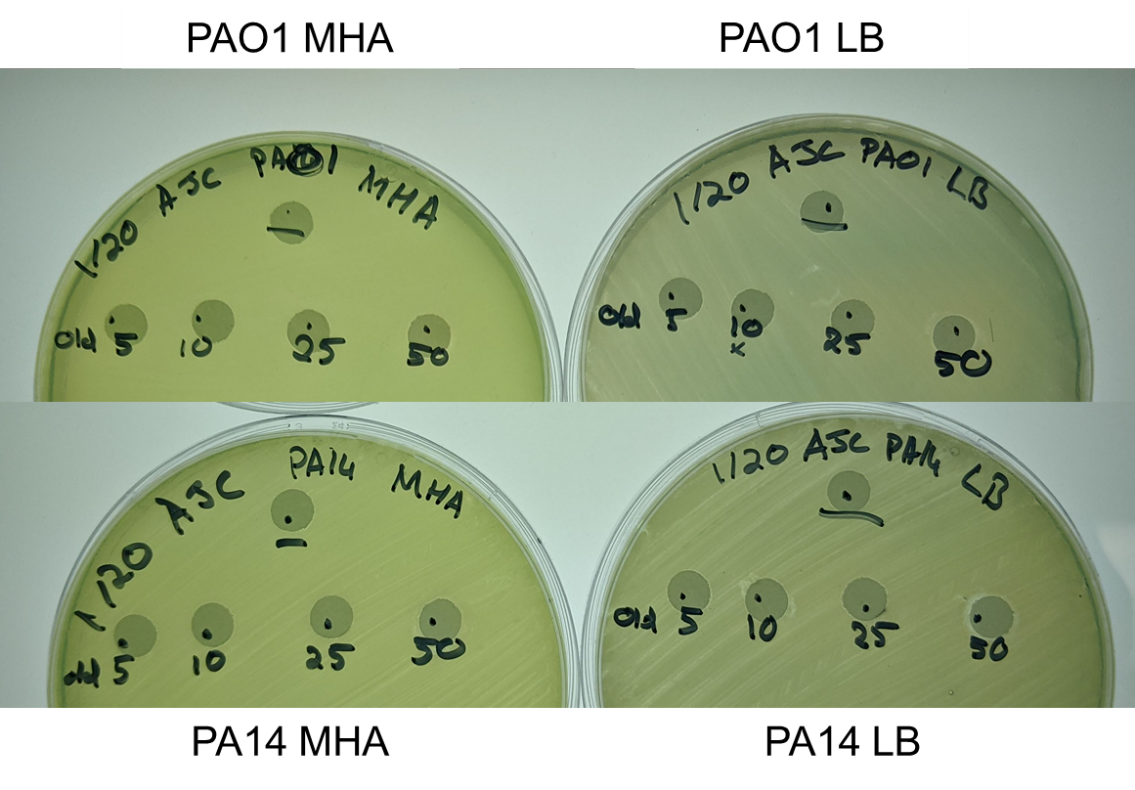

**Supp. Fig. 4** Assessment of inhibition of *P. aeruginosa* growth by manuka honey through a zone of inhibition assay. Discs were soaked in 5, 10, 25, or 50 v/v% manuka honey. “-” represents a filter disc without honey. Dry honey-containing filter disks were incubated on a lawn of *P. aeruginosa* (PA14 or PAO1) on either LB (lysogeny broth) or MH (Mueller-Hinton) agar plates for 16 hours. Plates were then imaged to demonstrate the extent of growth inhibition. There were no zones of inhibition present on any of the discs. Discs are 6 mm in diameter.

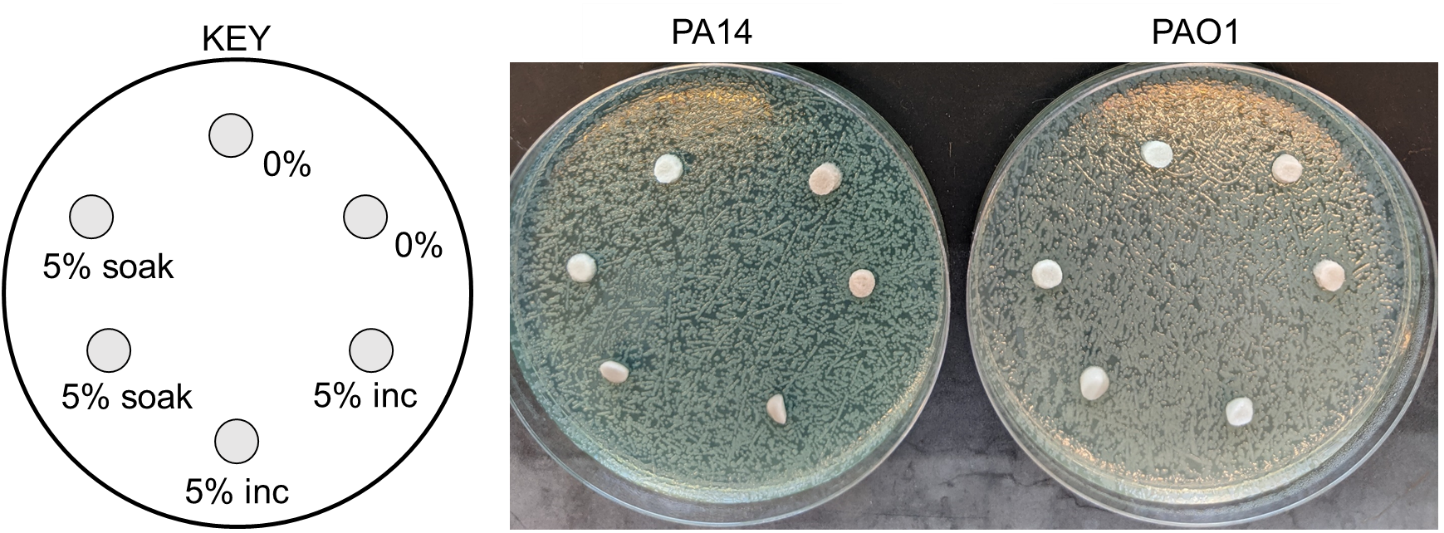

**Supp. Fig. 5** Zone of inhibition of hydrated scaffolds on lawns of PA14 or PAO1. Mineralized collagen scaffolds with 0% honey, 5% honey incorporated into the scaffold (5% inc), and 5% soaked into the scaffolds were added to lawns of *P. aeruginosa*. No zones of inhibition were visible. Scaffolds are 6 mm in diameter.

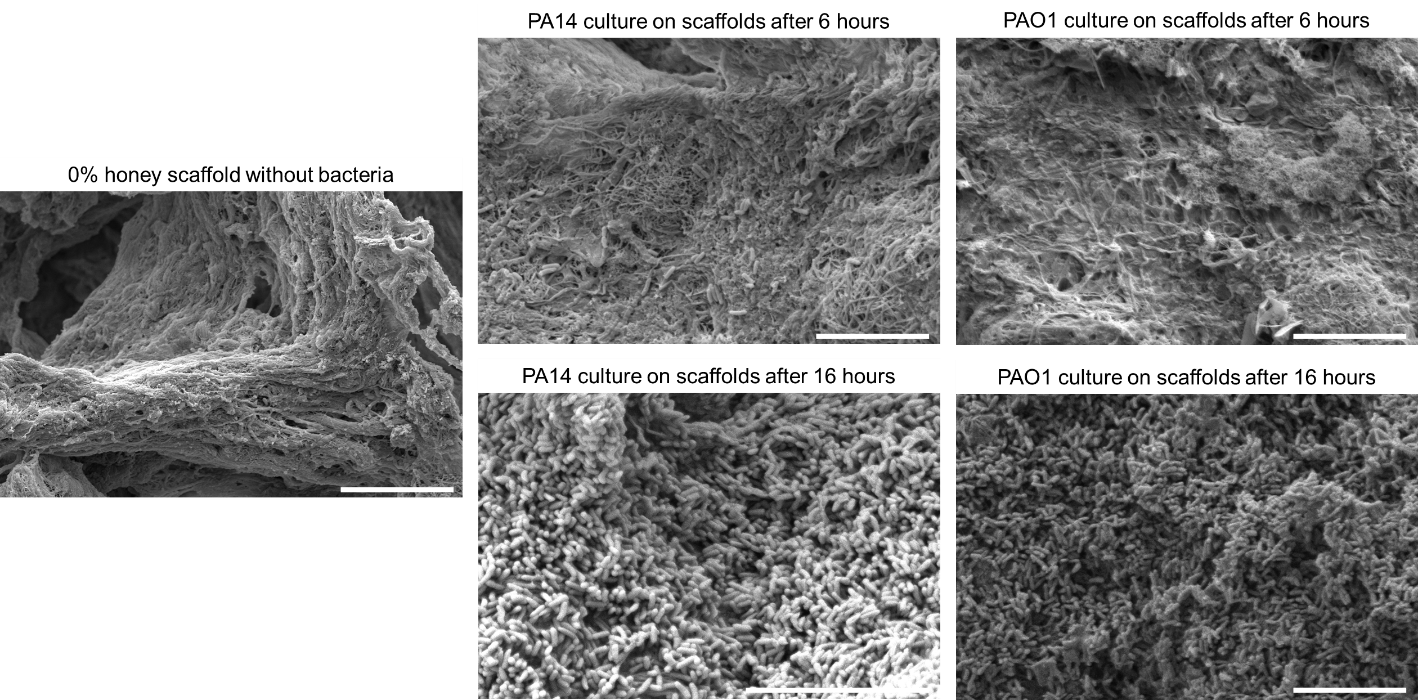

**Supp. Fig. 6** SEM images of 0% honey-containing scaffolds after 6 and 16 hours of PA14 and PAO1 strains of *Pseudomonas aeruginosa* culture in KA broth surrounding scaffolds. Scale bar represents 10 µm. Bacteria are adhered to the surface of scaffolds regardless of time and strain. PA14 bacteria uniformly coat the surface of the scaffold after 16 hours with minimal gaps.

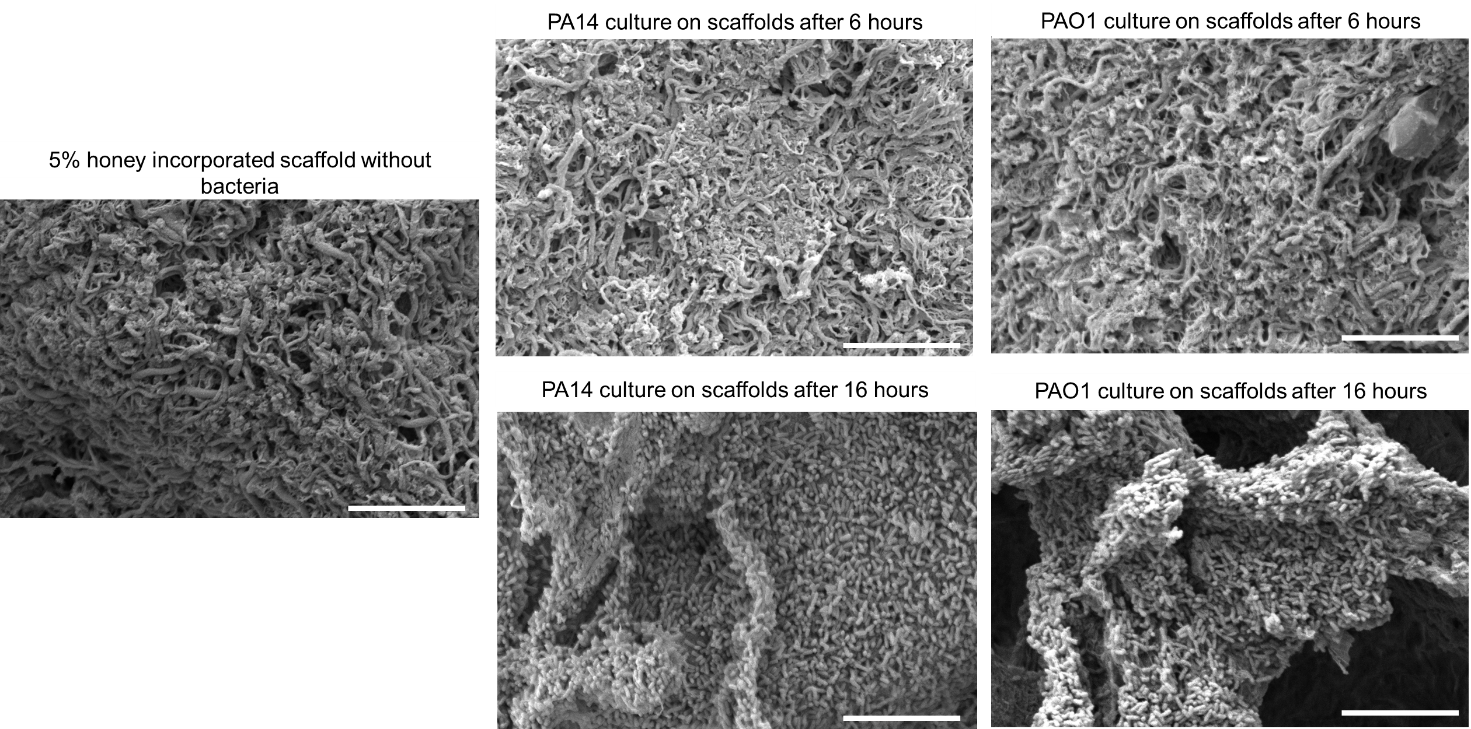

**Supp. Fig. 7** SEM images of 5% honey incorporated scaffolds after 6 and 16 hours of PA14 and PAO1 strains of *Pseudomonas aeruginosa* culture in KA broth surrounding scaffolds. Scale bar represents 10 µm. Bacteria are adhered to the surface of scaffolds regardless of time and strain. PA14 and PAO1 bacteria uniformly coat the surface of the scaffold after 16 hours with minimal gaps.

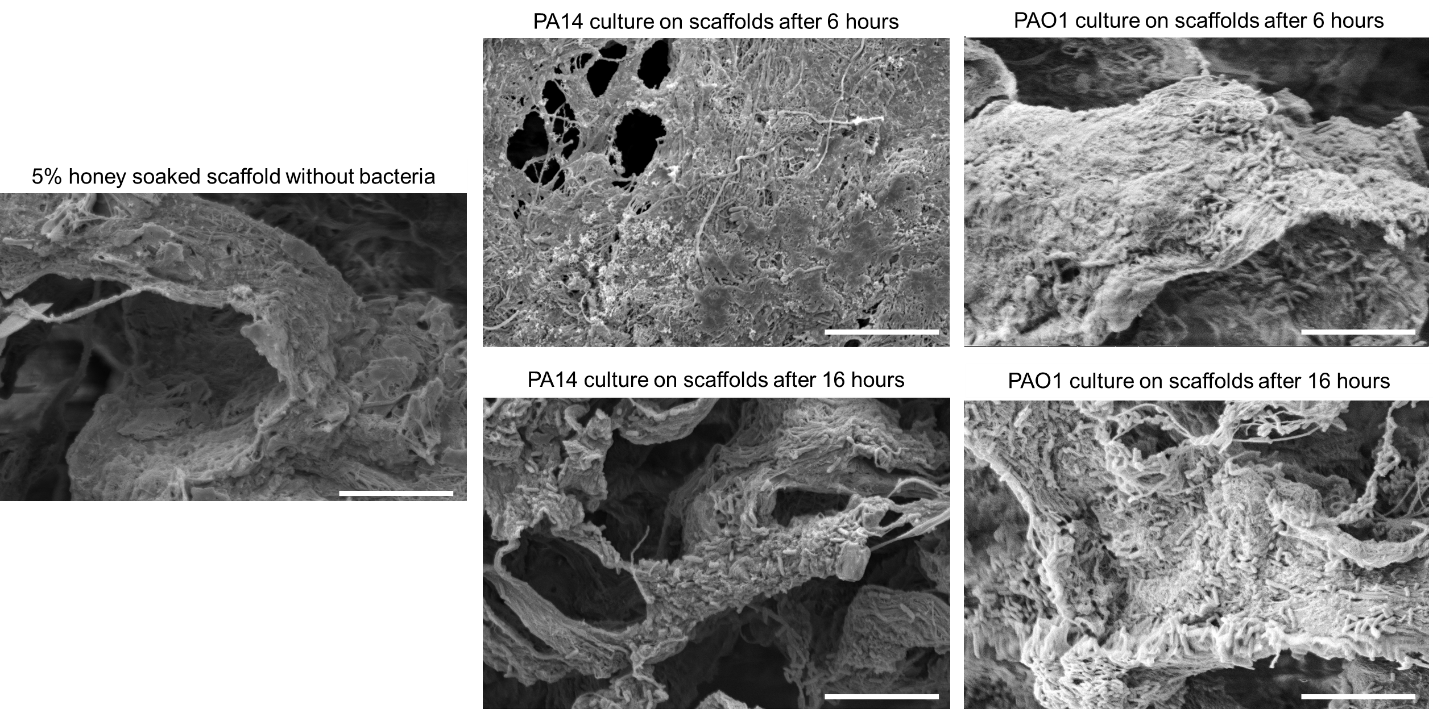

**Supp. Fig. 8** SEM images of 5% honey soaked scaffolds after 6 and 16 hours of PA14 and PAO1 strains of *Pseudomonas aeruginosa* culture in KA broth surrounding scaffolds. Scale bar represents 10 µm. Bacteria are adhered to the surface of scaffolds regardless of time and strain, but do not uniformly coat the surface after 16 hours.

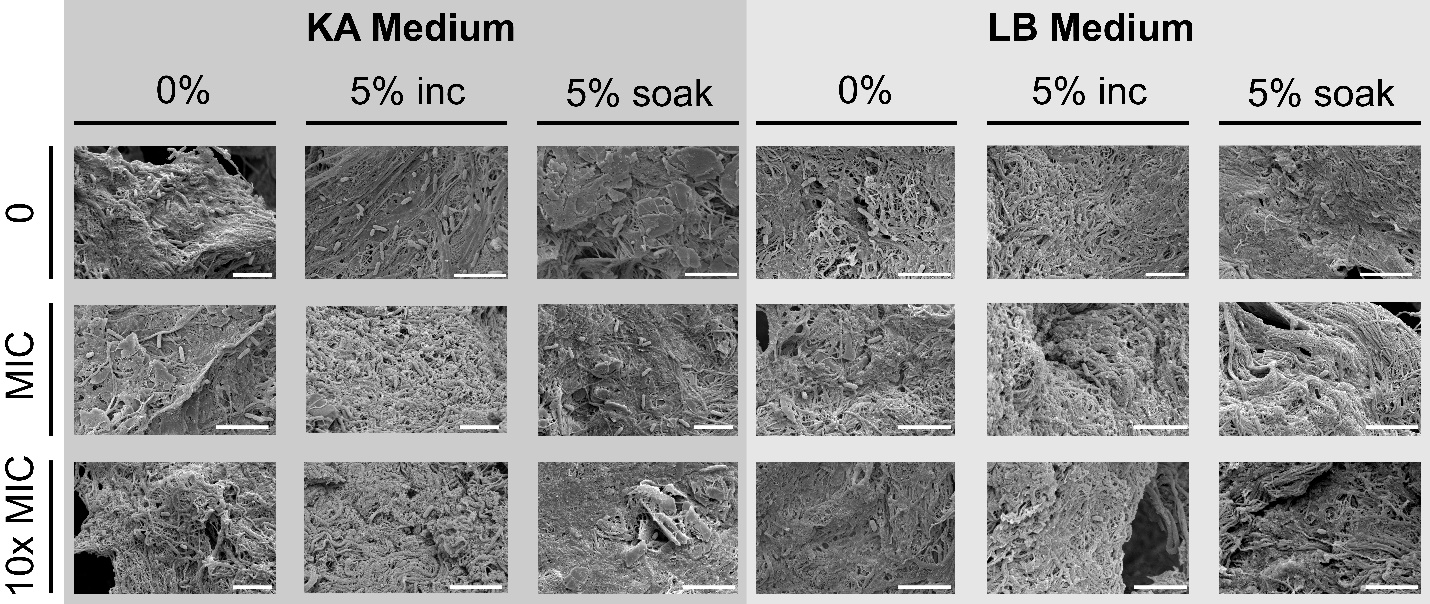

**Supp. Fig. 9** SEM images of *P. aeruginosa* (PA14) culture after 6hrs on mineralized collagen scaffolds with 0% honey (0%), 5% incorporated honey (5% inc), and 5% soaked honey (5% soak) with no additional antibiotics (0), minimum inhibitory concentration (MIC) with gentamicin, and 10 times the MIC (10x MIC). Scaffolds were cultured in KA medium and lysogeny broth (LB) medium. Scale bar represents 5 µm.
